# Alternative splicing in pediatric central nervous system tumors highlights oncofetal candidate *CLK1* exon 4

**DOI:** 10.1101/2024.08.03.606419

**Authors:** Ammar S. Naqvi, Patricia J. Sullivan, Ryan J. Corbett, Priyanka Sehgal, Karina L. Conkrite, Komal S. Rathi, Brian M. Ennis, Katharina E Hayer, Bo Zhang, Miguel A. Brown, Daniel P. Miller, Alex Sickler, Adam A. Kraya, Kaleem L. Coleman, Joseph M. Dybas, Zhuangzhuang Geng, Christopher Blackden, Shehbeel Arif, Antonia Chroni, Aditya Lahiri, Madison L. Hollawell, Phillip B. Storm, Dalia Haydar, Jessica B. Foster, Mateusz Koptyra, Peter J. Madsen, Sharon J. Diskin, Andrei Thomas-Tikhonenko, Adam C. Resnick, Jo Lynne Rokita

## Abstract

**Background:** Pediatric brain tumors are the leading cause of disease-related mortality in children, yet many aggressive tumors lack effective therapies. RNA splicing is a hallmark of cancer, but it has not yet been systematically studied in pediatric brain tumors.

**Methods:** We analyzed 729 pediatric brain tumors spanning histologies and molecular subtypes to quantify differential tumor splicing. We developed the *Splicing Burden Index (SBI)* to enable cross-sample comparisons and performed hierarchical clustering of highly variable splice events to define splicing-informed tumor groups. These were integrated with clinical outcomes, pathway activity, and proteogenomic data. Recurrent splice events were prioritized for predicted functional impact, and *in vitro* perturbation studies were performed targeting the splicing kinase *CDC-like kinase 1 (CLK1)*.

**Results:** SBI revealed substantial inter- and intra-histology heterogeneity. Clusters were enriched for histologies and molecular subtypes, several of which were independently associated with survival beyond histology and clinical covariates. Spliceosome pathway activity varied across clusters and was associated with worse survival, yet was not correlated with SBI, indicating distinct dimensions of splicing dysregulation. Functional prioritization identified a recurrent in *CLK1* exon 4, required for canonical kinase activity. *CLK1* exon 4 inclusion followed an oncofetal pattern and showed context-dependent associations with outcome distinct from total *CLK1* expression. Pharmacologic inhibition and exon 4-specific perturbation of *CLK1* reduced tumor cell viability and disrupted cancer-relevant splicing and transcriptional programs.

**Conclusions:** This study systematically characterizes splicing in pediatric brain tumors, identifies splicing-informed subgroups, and prioritizes *CLK1* exon 4 as an oncofetal tumor-specific event, motivating further preclinical exploration.

**Key Points:**

● Splicing analysis of 729 pediatric CNS tumors identifies splicing-defined clusters.
● *CLK1* exon 4 inclusion is widespread and developmentally regulated.
● Exon-level *CLK1* regulation shows context-dependent links to prognosis in aggressive CNS tumors.

## Importance of study

This study provides a comprehensive, cross-histology characterization of alternative splicing across 729 primary pediatric central nervous system (CNS) tumors, revealing widespread and heterogeneous splicing dysregulation with clinical relevance. By introducing the Splicing Burden Index and defining splicing-informed tumor clusters, we show that splicing patterns capture biologically and prognostically meaningful heterogeneity beyond traditional histologic classification. We further demonstrate that global splicing burden, spliceosome pathway activity, and exon-level regulation represent distinct dimensions of splicing biology. Functional prioritization of recurrent splice events identifies *CDC-like kinase 1 (CLK1)* exon 4 as a developmentally regulated, widely included splice event that produces the active kinase isoform and shows context-dependent associations with outcome. Experimental perturbation of *CLK1* exon 4 reduces tumor cell viability and disrupts cancer-relevant transcriptional and splicing programs. Together, these findings establish alternative splicing as a fundamental and targetable feature of pediatric CNS tumor biology and provide a framework for splicing-directed therapeutic strategies in high-risk disease contexts.

## Lay Summary

Brain tumors are the leading cause of cancer-related death in children, and many pediatric brain tumors remain difficult to treat. While most research has focused on genetic mutations, another important layer of regulation—how genes are “spliced” to make different versions of RNA and proteins—has been less well studied in childhood brain cancers. In this study, we analyzed RNA data from 729 pediatric brain tumors to understand how RNA splicing differs across tumor types. We found that tumors could be grouped into distinct categories based on their splicing patterns, and that these patterns were linked to differences in tumor behavior and patient outcomes. Some tumors showed widespread splicing changes that affected important cancer-related pathways. We also identified a splicing change in a gene called *CLK1* that is common in pediatric brain tumors and normally seen during early brain development. Changing how *CLK1* is spliced reduced tumor cell growth and disrupted cancer-related pathways in a representative patient-derived model. Importantly, drugs that target this pathway are already being tested in adults with cancer. Together, these findings show that RNA splicing plays a major role in pediatric brain tumor biology and may represent a new way to classify tumors and develop targeted treatments for children with brain cancer.

## Introduction

Pediatric brain cancer is the number one cause of disease-related death in children^1^. Despite advances in genomic profiling, many aggressive pediatric central nervous system (CNS) tumors lack effective targeted therapies, underscoring the need to identify additional regulatory mechanisms that drive tumor behavior and therapeutic vulnerability. One such mechanism is alternative pre-mRNA splicing, a fundamental post-transcriptional process that expands proteomic diversity and fine-tunes gene regulation, yet remains incompletely characterized in pediatric CNS tumors.

Prior studies have demonstrated that splicing alterations can contribute to pediatric brain tumor biology through diverse mechanisms. Rare, mutually exclusive mutations in spliceosome-associated factors such as *SF3B1* and *SF3B2* have been identified in pediatric high-grade gliomas (HGGs), disrupting processes involved in DNA replication, genome integrity, or transcriptional fidelity^2^. More recent work has shown that alternative splicing can activate oncogenic pathways in pediatric HGGs, including RAS/MAPK signaling, and is associated with worse clinical outcomes^3^. Functional relevance of splice isoforms has also been demonstrated *in vivo*, where a tumor-specific isoform of the neuronal cell adhesion molecule *NRCAM*, but not its canonical transcript, was essential for pediatric HGG xenograft growth^4^. Additional studies have implicated *OTX2*-driven splicing programs in Group 3 medulloblastoma (MB)^5^, U1 snRNA splice site mutations in SHH MB^6^, tumor-restricted splice-derived neoantigens in MB^7^, and pathogenic *NF1* splice variants in Neurofibromatosis Type I–associated tumors^8^.

Collectively, these studies establish alternative splicing as a biologically important contributor to pediatric brain tumor pathogenesis. However, they have largely focused on isolated mechanisms, specific tumor types, or individual splice events. As a result, the broader architecture of splicing dysregulation across pediatric CNS tumors—and how exon-level splicing integrates with tumor lineage, signaling pathways, and clinical outcome—remains poorly defined. Particularly, it is unclear which splicing alterations are merely correlative and which actively rewire protein function, signaling networks, or immune visibility to drive malignant behavior.

Alternative splicing is especially relevant in the developing brain, which exhibits the most complex and conserved splicing programs of any tissue^9,10^. These programs are tightly regulated by *trans*-acting RNA-binding proteins (RBPs), including Serine-rich Splicing Factors (SRSFs) and heterogeneous nuclear ribonucleoproteins (hnRNPs), whose dysregulation can profoundly alter cellular identity and function^11,12^. Perturbation of these regulatory networks, or of kinases that modulate splicing factor activity, has the potential to create lineage-specific dependencies in cancer.

Here, we perform a comprehensive, cross-histology analysis of transcriptome-wide alternative splicing across 729 pediatric CNS tumors spanning diverse histologic and molecular subtypes. We identify widespread alternative splicing alterations that define splicing-informed tumor subgroups, reveal heterogeneity within and across tumor lineages, and uncover recurrent splice events predicted to alter protein function. Using a functional prioritization framework, we identify CDC-like kinase 1 (*CLK1*) as a candidate regulator of oncogenic splicing programs in pediatric CNS tumors. Canonical *CLK1* activity requires inclusion of exon 4, whereas skipping of this exon produces catalytically inactive isoforms^13^. Prior studies have identified *CLK1* as a therapeutic target in multiple adult cancers^14,15,16–20^, but none have examined its role in pediatric cancer. Here, we demonstrate that exon 4 inclusion in *CLK1* is widespread, developmentally-regulated, and functionally relevant in pediatric CNS tumors, supporting the concept that splice-defined states may represent actionable layers of tumor biology.

## Materials and Methods

### Study participants

Study participants include pediatric brain tumor patients whose genomic data were deposited into and obtained from the OpenPedCan^21^ project. Histologies include atypical teratoid rhabdoid tumor (ATRT, N = 24), choroid plexus tumor (CPT, N = 20), craniopharyngioma (CPG, N = 29), diffuse midline glioma (DMG, N = 29), ependymoma (N = 69), germ cell tumor (N = 8), low-grade glioma (LGG, N = 200), medulloblastoma (MB, N = 105), meningioma (N = 17), mesenchymal tumor (N = 17), mixed neuronal-glial tumor (GNT, N = 63), neurofibroma plexiform (N = 10), non-neoplastic tumor (N = 26), other CNS embryonal tumor (N = 8), other high-grade glioma (other HGG, N = 74), schwannoma (N = 13), and Rare CNS tumors (N = 17) (**Table S1**).

### Primary data analyses

Somatic primary workflows were implemented by the Kids First Data Resource Center as described in the Open Pediatric Brain Tumor Atlas (OpenPBTA)^22^ and OpenPedCan[Citation error] projects. The code for these workflows, including RNA-seq quantification, fusion identification, RNA splicing, and SNV, INDEL, CNV, SV calling, can be found at https://github.com/d3b-center/OpenPedCan-workflows. Sample-level data can be found through the Kids First Portal at https://kidsfirstdrc.org/. To detect alternative splicing, we ran rMATS turbo (v. 4.1.0)^23^ with GENCODE v39 GFF annotations on single samples, as described by the Kids First RNA-Seq workflow (https://github.com/d3b-center/OpenPedCan-workflows). We filtered for alternative splicing events with ≥ 10 junction read counts. To compare RNA-Seq from *CLK1* exon 4 morpholino-treated cells vs control morpholino-treated cells, we ran rMATs with three biological replicates for each condition ‘--b1 --b2’. This grouped analysis calculated ΔPSI, p-values, and FDR statistics for each splice event. These results were then used for all downstream processing throughout the manuscript.

### Cell Culture

The high-grade glioma patient-derived cell lines 7316-1763 and 7316-1769 were obtained by CBTN request, and the KNS-42 cell line was obtained from Accegen (ABC-TC0532). The pediatric HGG cell line KNS-42 was cultured in DMEM-F12 (GIBCO, 11320033) supplemented with 10% FBS (GIBCO, 26140079), 2 mmol/L L-glutamine (GIBCO, 25030081), and 1X penicillin/streptomycin (GIBCO, 15140122) at 37°C and 5% CO_2_. The cell line was authenticated by Guardian Forensic Sciences (Abington, PA) using the GenePrint 24 (Promega, B1870) short tandem repeat kit. Cells tested negative for mycoplasma using the EZ-PCR Mycoplasma Detection Kit (Biological Industries, 20-700-20) and were used for a maximum of 12 passages post thaw.

### Morpholino Treatments

A Vivo-Morpholino ACTCTTCTGGAAACGTCAAGTGGGC (Gene Tools, LLC) targeting the intron 3-exon 4 splice junction was used to skip exon 4 in *CLK1*. Cells were treated with 1, 5, and 10 μM concentrations of *CLK1* morpholino and 10 μM of Control morpholino. 48 hours post-treatment, cells were harvested for PCR and immunoblots.

### RNA Extraction and Quantitative Real-time PCR (qRT-PCR)

Total RNA was isolated and treated with DNAse using the Maxwell RSC simplyRNA Cells kit (Promega, AS1390) with the Maxwell RSC48 Instrument (Promega) per the manufacturer’s instructions. Next, 2 μg of RNA were reverse-transcribed using SuperScript IV (Invitrogen, 18090010). Primers used for *CLK1* mRNA transcript quantification are listed in **Table S6K**. qRT-PCR was performed using PowerSYBR Green PCR Master Mix (Invitrogen, 4367659) on an Applied Biosystems Viia7 machine. The amplification was performed using the following settings: denaturation at 95°C for 10 min, followed by 40 cycles of denaturation at 95°C for 15 s and annealing at 60°C for 1 min. The comparative cycle threshold (CT) method was applied to quantify the expression levels of *CLK1*. The fold change of gene expression was calculated by the equation 2ΔΔCT, with *HPRT* (Thermo Fisher, 4453320, assay ID: Hs02800695_m1) used as the housekeeping gene.

### Protein Extraction

Cultured cells were washed once in chilled D-PBS (pH 7.4) and lysed in RIPA buffer containing 50 mM Tris-HCl, pH 7.4, NP-40 (1%), deoxycholate (0.25%), 150 mM NaCl, 1 mM EDTA pH 8.0, 1x protease and phosphatase inhibitor cocktail (Pierce Halt Inhibitor Cocktail, Thermo Fisher Scientific, 78446), and SDS (0.1%). Total protein in the lysate was estimated by the DC Protein assay (BioRad Laboratories, 5000111).

### Detection of Proteins Using Immunoblot Analysis

70 μg of total protein were mixed with 5X SDS loading dye (Biorad, 161-0374) and resolved on 10% SDS-polyacrylamide gel. The protein was transferred onto a PVDF membrane (Immobilin-P, Millipore, IPVH00010) and probed with α-CLK1 mouse monoclonal primary antibody (Santa Cruz, sc-515897) and HRP-conjugated secondary antibody (Cell Signaling Technology, 7076S). Bands were detected using enhanced chemiluminescence (Millipore, WBKLS0500) and captured by a Chemiluminescence imager (GE Healthcare). β-actin was used as the loading control and probed with α-β-actin rabbit monoclonal antibody (Cell Signaling Technology, 12262S).

### pan-DYRK/CLK1 inhibitor Cirtuvivint (SM08502) experiments

The KNS-42 cell line was cultured in DMEM-F12 (Gibco, 11330032) supplemented with 10% FBS (Corning, MT3501CV, lot 003322001) and additional L-glutamine (Thermo Fisher, 25030081) to a final concentration of 4.5 mM. Dissociation was performed with Trypsin-EDTA (0.05%, Thermo 25300054) and counted on a DeNovix Cell Drop cell counter.

For growth kinetics, 10,000 (3 day assay) or 6,000 (6 day assay) cells were plated per well into a 96-well plate (Greiner Bio-One, 655098) in a 200 uL total volume per well. Plates were placed into an Incucyte SX5 device and scanned every 2 hours for several days to measure growth via a mask designed uniquely for this cell type. At the end point of the assay, cell viability was analyzed with CellTiter Glo 2.0 reagent (Promega, G9242) by replacing half the media with reagent and reading on a Promega GloMax device.

Cirtuvivint (MedChem Express, HY-137435) was resuspended in 100% DMSO (Sigma, D2650-5X5ML) to 1 mM and stored in aliquots at -80 C. Dosing was optimized via serial dilution at a range of 20 uM to 0.02 uM against a vehicle control equivalent to the highest dosing of drug. Cells were plated and at 24 hours, 100 uL of media were removed from each well and replaced with drug media for a final dose range of 0.01, 0.05, 0.5, 0.1, 1, 5, and 10 uM. Cells were untouched for 3 days total while growth was monitored via Incucyte.

### Cell Viability Assay

Cell viability was measured using the CellTiter-Glo (CTG) luminescent cell viability assay (Promega, G7570). Cells were seeded in white 96-well flat-bottom plates at a density of 24,000 cells per well and treated the following day with either 7.5 μM control or CLK1 exon 4 targeted morpholino. Luminescence was measured using a Biotek Synergy 2 plate reader at 24, 48, 72, and 96 hours.

## Results

### Pediatric brain tumors display heterogeneous global patterns of differential splicing

To investigate alternative splicing in pediatric brain tumors, we analyzed total RNA-Seq data from 729 diagnostic tumors from the Open Pediatric Cancer (OpenPedCan) project^21^ (**Figure 1A**). Demographic and clinical characteristics for each patient and tumor are provided in **Table S1**. We quantified splicing at single-sample resolution using percent-spliced-in (PSI) values computed with rMATS turbo^23^, which measure relative exon or junction usage independent of total gene expression, and identified skipped exon (SE), alternative 5’ splice site (A5SS), alternative 3’ splice site (A3SS), and retained intron (RI) events.

**Figure 1:**
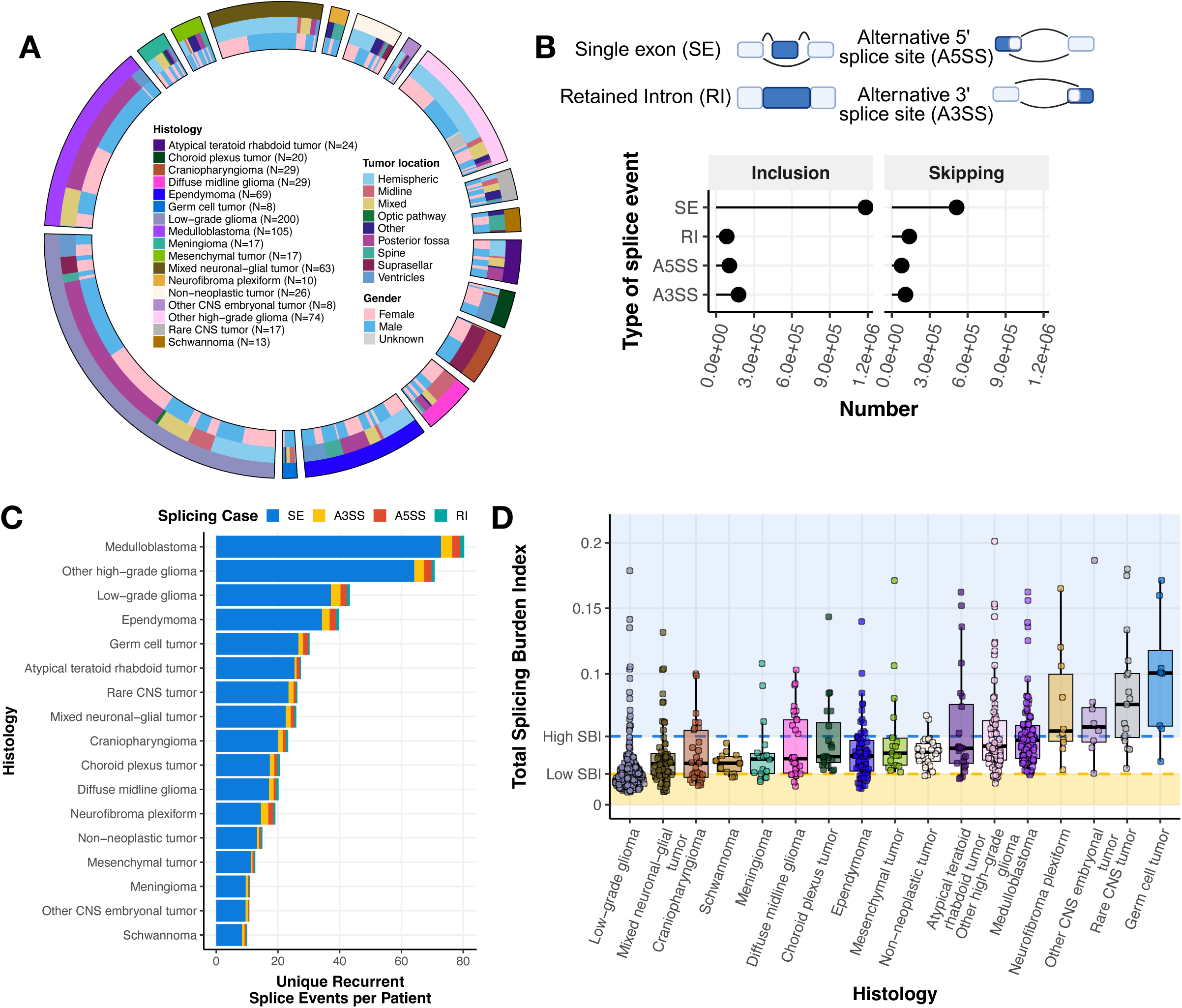
Pediatric brain tumors display heterogeneous global patterns of differential splicing. **(A)** Circos plot of CNS tumors used in this study, categorized by histology, tumor location, and reported gender. Non-neoplastic tumors consist of benign tumors and/or cysts. Rare tumors are CNS tumors with a small (<4) number of patient samples. **(B)** Lollipop plot illustrating the total number of recurrent differential splicing events across the cohort, classified by splicing type (SE: single exon, RI: retained intron, A5SS: alternative 5’ splice site, A3SS: alternative 3’ splice site, top panel created with BioRender.com). **(C)** Barplots of the number of histology-specific recurrent SE splicing events per patient. Histologies are reverse-ordered by the total number of unique events (skipping + inclusion). **(D)** Distribution plots of splicing burden index (SBI) of all splicing events and types by histology. Shaded regions represent high (blue; ≥ Quartile 3) and low (yellow; ≤ Quartile 1) SBI groups.

Given the biological and molecular diversity of pediatric CNS tumors, we asked whether distinct splicing patterns would be observed across histologies and molecular subtypes. Among recurrent (N ≥ 2) differential splice events, SE events were the most frequent (**Figure 1B**), consistent with a prior report in pediatric HGGs^3^. We next assessed whether these events were histology-specific or shared across tumor types. We observed both shared and histology-specific splicing events (**Figure S1A**), with medulloblastoma (MB), low-grade glioma (LGG), and non-DMG high-grade glioma (other HGG) tumors exhibiting the largest number of unique recurrent events, consistent with their larger representation in the cohort. Importantly, this pattern persisted after normalizing for the number of patients within each histology (**Figure 1C**), indicating that both shared and histology-enriched splicing programs are present across pediatric CNS tumors. A complete list of unique events per histology is reported in **Table S2A**.

To enable cross-sample comparisons of alternative splicing activity, we developed the splicing burden index (SBI): a sample-level metric quantifying the proportion of differential alternative splicing events in each tumor (**Online Methods**). Across the cohort, the median SBI was 0.0351 (3.51%). Among tumor types, LGGs exhibited the lowest median SBI (2.16%), while germ cell tumors (GCTs) had the highest (10.1%) (**Figure 1D**). Notably, SBI varied substantially within tumor histologies. Tumor groups with high SBI variance (> 3rd quartile variance) included other CNS embryonal tumors, GCTs, rare CNS tumors, neurofibroma plexiforms, and ATRTs, indicating marked inter-tumoral heterogeneity in splicing programs. Although overall SBI values were modest on a cohort-wide scale, this variability underscores the heterogeneity of splicing programs across pediatric CNS tumors.

We next asked whether tumors with a low tumor mutation burden (TMB) might exhibit an increased splicing burden as an alternate mechanism contributing to tumorigenesis. Instead, we observed a very weak positive correlation between TMB and SBI (Pearson’s R = 0.13, p-value = 6.9e-4; **Figure S1B**), which persisted after excluding hyper-mutant tumors (R = 0.14, p-value = 3.4e-4, **Figure S1C**). Although statistically significant, these correlations explain little variance, indicating that TMB and splicing burden are largely independent across the cohort. Taken together, these results suggest that TMB is not a major determinant of global splicing burden in pediatric CNS tumors. When stratified by histology, significant positive correlations were observed only in GCTs (R = 0.79, p = 0.033) and MB (R = 0.21, p = 0.048), whereas schwannomas showed a significant inverse relationship between TMB and SBI (R = -0.78, p = 0.0049) (**Figure S1D**). Schwannomas also exhibited the lowest variance in SBI across tumor types, consistent with tighter constraint on splicing burden in this lineage.

### Splice events cluster pediatric brain tumor histologies and are associated with survival outcomes

To assess whether CNS tumors exhibit shared patterns of alternative splicing, we performed hierarchical clustering of samples using PSI values from the top 5,000 most variable splice events across all primary tumors (N = 729). This analysis identified 10 distinct clusters, each enriched for specific histologies and/or molecular subtypes (**Figure 2A-B**, and **S2A-D**). Notable enrichments included Cluster 4 with ependymoma (OR = 222.8), Cluster 1 with LGGs (OR = 47.9), and Cluster 9 with MB (OR = 115.1), with the latter driven predominantly by Group 4 tumors (OR = 94.4) (**Figure S2B**). These results are consistent with prior work demonstrating that the MB subgroups WNT, SHH, Group 3, and Group 4 can be delineated based on splicing patterns inferred by expression arrays^24^. HGGs, including DMGs, exhibited the greatest splicing heterogeneity, with samples distributed across 8 of the 10 clusters (**Figure S2C-D**). Cluster membership for all samples is provided in **Table S2B**.

**Figure 2:**
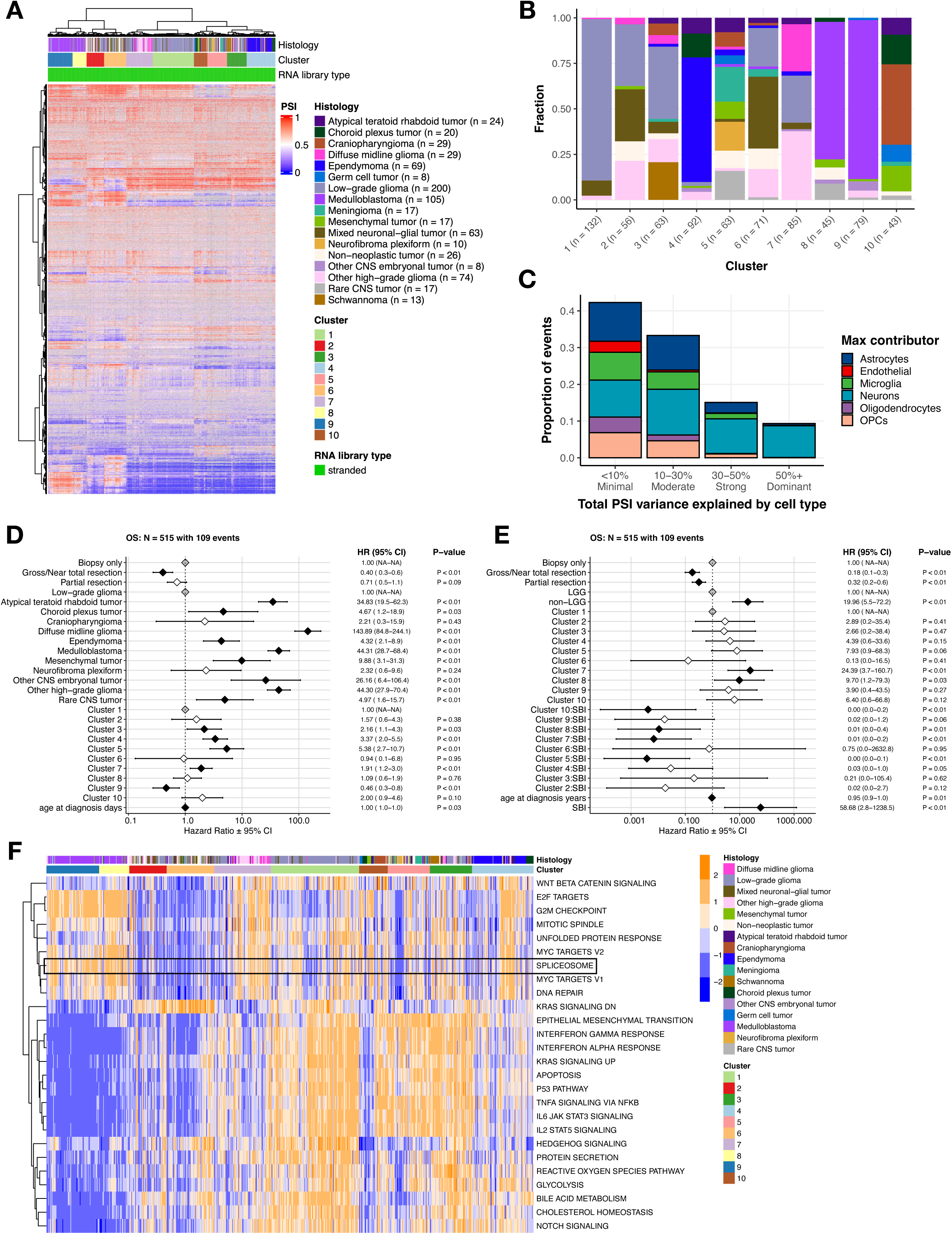
Splicing-based clustering reveals heterogeneous tumor groups with distinct histologies, pathway activities, and clinical outcomes. **(A)** Hierarchical clustering heatmap of PSI values for top 5,000 most variable splice events across tumors. **(B)** Stacked barplot showing histology sample membership fraction for each cluster. **(C)** Stacked barplot showing the contribution of PSI variance for each cell-type composition. **(D)** Forest plot of cox proportional hazards multivariate interaction OS model for clusters relative to Cluster 1, including covariates for tumor resection and age at diagnosis. Black and white diamonds indicate statistically significant and not significant hazard ratios (HRs), respectively, with intervals denoting 95% confidence intervals. Gray diamonds indicate reference levels of factor covariates. **(E)** Forest plot of cox proportional hazards multivariate interaction OS model for clusters, including covariates for tumor resection, age at diagnosis, and cluster SBI. Black and white diamonds indicate statistically significant and not significant hazard ratios (HRs), respectively, with intervals denoting 95% confidence intervals. Gray diamonds indicate reference levels of factor covariates. **(F)** Heatmap of top cancer-related pathways by cluster (GSVA scores represented by blue/orange color).

To assess whether these clustering patterns primarily reflected differences in cellular composition rather than tumor-intrinsic splicing programs, we quantified the proportion of PSI variance explained by estimated cell-type composition inferred from bulk RNA-Seq gene expression data using BRETIGEA^25^. For most cluster-defining events, less than 30% of total PSI variance was attributable to cell-type composition; approximately 15% of events showed 30–50% variance explained, and 10% exceeded 50% (**Figure 2C**). These findings indicate that while cellular composition contributes to splicing variability for a subset of events, it does not account for the majority of splicing differences that define the observed clusters.

We next examined overall survival (OS) across splicing-defined clusters. After adjusting for histology and additional clinical covariates including tumor resection and age, several clusters remained independently associated with OS relative to Cluster 1 (**Figure 2D**). Specifically, Cluster 9 showed improved OS, whereas Clusters 3, 4, 5, and 7 were associated with worse OS. These results indicate that splicing-defined clusters are associated with prognostically relevant heterogeneity beyond histologic classification alone. Given these associations, we evaluated SBI as a covariate in the survival models and tested for interactions between SBI and cluster membership. We identified significant SBI-cluster interactions in Clusters 5, 7, 8, and 10, indicating that the relationship between splicing burden and OS varies across splicing-defined contexts. In Clusters 5, 7, 8, and 10, higher SBI was associated with improved OS within those clusters. However, at the cluster level, membership in Clusters 7 and 8, as well as higher SBI values across the cohort, remained associated with poorer OS (**Figure 2E**). SBI distributions by cluster, stratified by histology, are shown in **Figure S2E**.

To explore the functional impact of splicing alterations, we identified differentially-expressed cancer-associated signaling pathways across the 10 splicing-defined clusters using GSVA (**Figure 2F**). The KEGG spliceosome pathway was significantly upregulated in Clusters 7, 8, and 9, which were associated with poorer OS (**Table S2C,** Bonf-adj p < 0.05; **Figure S3A**) and significantly downregulated in Clusters 2, 5, and 6, which were associated with more favorable outcomes (**Table S2,** Bonf-adj p < 0.05; **Figure S3A**). These results indicate marked heterogeneity in spliceosome pathway activity across splicing-defined tumor contexts.

We next assessed whether spliceosome activity, quantified by GSVA score, was associated with survival outcomes across the cohort using a multivariate cox proportional hazards model adjusting for tumor resection, glioma grade group, age at diagnosis, and cluster membership. Tumor resection was a favorable prognostic factor (HR = 0.25 for gross/near total and 0.44 for partial, p < 0.01), whereas non-LGG tumors (HR = 13.60, p < 0.01), Cluster 7 membership (HR = 4.36, p < 0.01), and higher spliceosome GSVA scores (HR = 2.91, p < 0.01) were independently associated with worse OS (**Figure S3B**).

To evaluate whether increased KEGG spliceosome GSVA scores reflect elevated protein abundance, we integrated RNA expression with matched proteogenomic data (N = 122) from the Clinical Proteomic Tumor Analysis Consortium (CPTAC)^26^. Protein (**Figure S3C**) and RNA (**Figure S3E**) expression of KEGG spliceosome pathway genes (**Table S3A**) were visualized alongside GSVA scores, revealing a significant positive correlation between protein abundance and GSVA scores (R = 0.4, p = 2.6e-6; **Figure S3E**), supporting the use of transcript-based GSVA as a proxy for pathway activity. Notably, spliceosome GSVA scores were not correlated with splicing burden (**Figure S3F**), indicating that global spliceosome pathway activity and splicing burden capture distinct aspects of splicing dysregulation.

### Widespread splicing alterations are associated with expression changes in splicing factors and recurrent splice events with predicted functional impact

To investigate potential mechanisms underlying widespread splicing alterations, we assessed recurrent somatic alterations across the cohort (N = 657, **Figure S4A**). While recurrent alterations were observed in canonical oncogenic drivers, recurrent predicted deleterious mutations in components of the splicing machinery were rare overall. Specifically, only 3.5% (23/657) of tumors harbored PolyPhen- and SIFT-predicted deleterious mutations in 50 of the 150 HUGO-annotated spliceosome genes (**Table S3B, S3D**) and 18.1% (119/657) harbored predicted deleterious mutations across 446 of the 1,350 known SF genes (**Tables S3C, S3E**). These findings indicate that widespread splicing alterations in pediatric CNS tumors are unlikely to be primarily driven by recurrent coding mutations in splicing machinery genes.

Prior studies have shown that in the absence of recurrent splicing factor mutations, changes in the expression of splicing regulators can influence downstream splicing programs and contribute to tumorigenesis^27–29^. Guided by this framework, we performed differential gene expression (DE) analysis between high and low SBI tumors, focusing on known splicing factors and related genes^30^ (**Table S3C**). We found 65.3% (N = 881/1350) of these genes were significantly differentially-expressed (adjusted p-value < 0.05, **Figure 3A-B, Table S3F**). Notably, 88% (30/34) of genes encoding the SRSFs and hnRNPs – families of *trans*-acting splicing regulators with established roles in exon-associated splicing^31^ – were significantly differentially expressed between high and low SBI tumors, suggesting that SBI reflects coordinated splicing factor–associated regulatory states.

**Figure 3:**
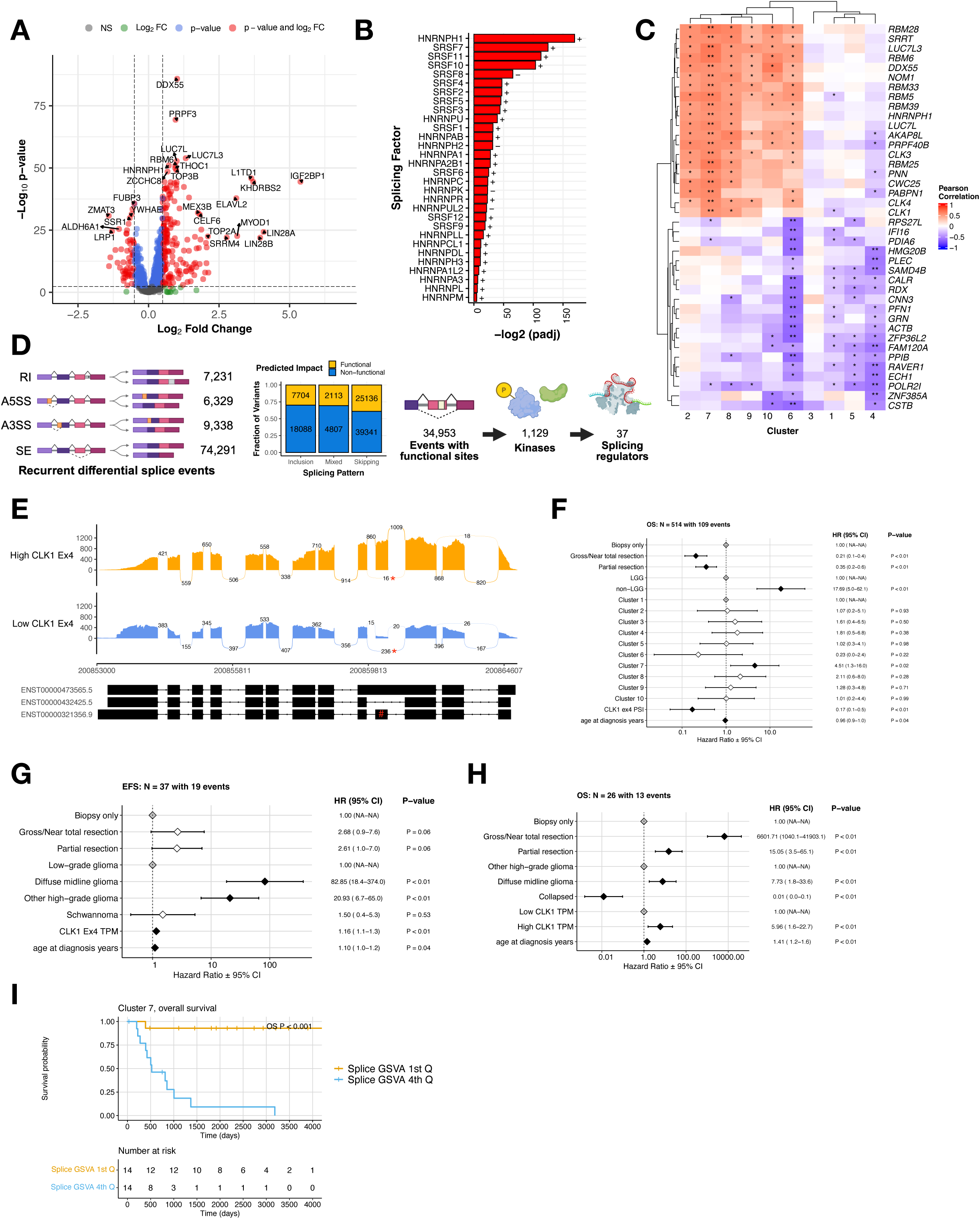
Splicing state-associated event prioritization highlights *CLK1* exon 4 as a recurrent functional splice event. **(A)** Volcano plot illustrating differentially-expressed splicing factor genes with high SBI compared to those with low SBI (NS = not significant, FC = fold change, colored dots represent log_2_FC > |.5| and/or Benjamini and Hochberg (B-H) adjusted p-value < 0.05) **(B)** Barplot showing significance of canonical trans-acting splicing factors. Fold-change directionality is annotated as + or -. **(C)** Correlation heatmap showing Pearson R between SBI and SF gene expression, stratified by cluster. Stars denote * = FDR < 0.05 and ** = FDR < 1.0e-5. **(D)** Workflow to prioritize candidate differential exon-level splicing events that alter UniProt-defined functional sites, created with BioRender.com. Stacked bar plots represent the fraction of exon inclusion, skipping, or mixed splicing events categorized by predicted impact. **(E)** Sashimi plot of two representative tumor samples with either high (BS_30VC3R2Q) or low (BS_FYP2B298) *CLK1* exon 4 inclusion. Reads supporting exon 4 skipping are marked with an asterisk (*), and exon 4 is indicated by a hash (#) on the transcript plot. **(F)** Cox proportional hazards model forest plot of OS for cluster membership, including covariates for extent of tumor resection, histology group, *CLK1* exon 4 PSI, and age at diagnosis. Black and white diamonds indicate statistically significant and not significant hazard ratios, respectively, with intervals denoting 95% confidence intervals. Gray diamonds indicate reference levels of factor covariates. **(G)** Cox proportional hazards model forest plot of EFS for Cluster 3 and **(H)** OS for cluster 7 adjusted for exon 4-included *CLK1* transcript abundance, including covariates for extent of tumor resection, histology group, and age at diagnosis. “Collapsed” indicates combined histologies with N < 3 to retain these tumors in the model. Black and white diamonds indicate statistically significant and not significant hazard ratios, respectively, with intervals denoting 95% confidence intervals. Gray diamonds indicate reference levels of factor covariates. **(I)** Kaplan–Meier curve for overall survival (OS) in Cluster 7, stratified by high vs low GSVA score.

To further assess whether global splicing burden reflects coordinated regulation of splicing machinery, we examined correlations between SBI and the expression of splicing factor (SF) genes across splicing-defined clusters. We generated a correlation heatmap of the top 40 SF genes most correlated with SBI, stratified by cluster (**Figure 3C**), with corresponding correlation statistics summarized by cluster and by histology (**Table S4A-B**). SBI showed significant positive correlations with SF gene expression in Clusters 2, 6, 7, 8, 9, and 10, whereas Clusters 1, 3, 4, and 5 exhibited predominantly negative correlations. Notably, Cluster 6 displayed a mixed pattern, with subsets of SF genes showing both positive and negative correlations with SBI. These findings indicate that increased splicing burden is associated with coordinated, but cluster-specific, regulation of splicing factor expression rather than uniform upregulation of the splicing machinery across all tumors.

To further prioritize recurrent splice changes with potential biological relevance across splicing-defined tumor states, we developed a robust and adaptable workflow to identify recurrent (N ≥ 2) differentially-spliced (ΔPSI z-score > |2|) events, ΔPSI > 0.2 (sample compared to cohort), and predicted functional impact (**Figure 3D**). Across the cohort, we identified 97,189 recurrent differential splicing events. Of these, 34,953 events were prioritized based on predicted gain or loss of annotated UniProt functional features, including disulfide bonds (N = 3,627), localization signals (N = 1,551), amino acid modification sites (N = 6,997), and protein domains (N = 28,779) (**Table S5**).

To focus on potentially targetable splice alterations, we restricted this set to functional splice events occurring in kinases, yielding 1,129 events. We further refined this list by prioritizing kinases with established roles in splicing regulation, resulting in 37 candidate splice events across 11 genes: *CLK1, CLK2, CLK3, CLK4, FASTK, MARK2, PKN2, PRKDC, SMG1, SRPK1,* and *SRPK3*. Applying additional filters based on expression (TPM > 10) and recurrence (> 10 tumors), we identified *CLK1, CLK3, FASTK,* and *PRKDC* as top candidates comprising 11 unique splice events. We excluded seven RI events, which are known to have higher false-positive rates in short-read sequencing^32^, and two additional RI events in *CLK3* that were annotated as A5SS and A3SS. After these exclusions, only two SE events remained: *CLK1* exon 4 and *PRKDC* exon 80. Comparison of *PRKDC* exon 80 inclusion across normal polyA-sequenced cohorts showed that our PBTA polyA data closely matched GTEx (**Figure S4B**), indicating that the stranded-specific signal may reflect a technical artifact. We therefore prioritized *CDC-like kinase 1 (CLK1)*, a well-established regulator of alternative splicing^11^, as a candidate node within high-burden splicing states for further investigation. *CLK1* regulates SRSF proteins through hyper-phosphorylation of SR-rich domains, promoting cooperative RNA binding and splicing activity^33–35^. Canonical *CLK1* activity requires inclusion of exon 4^13^. We observed differential splicing of this exon, with 34 tumors showing significant skipping and a mean ΔPSI of 0.29 (**Table S5C)**. This event is illustrated using sashimi plots contrasting tumors with high versus low exon 4 inclusion (**Figure 3E**). Notably, the majority of tumors (n = 695 out of 729) showed high *CLK1* exon 4 inclusion (mean PSI = 0.790; **Table S5D**); **Figure S4C**), motivating further investigation of this splice event in the context of tumor biology and development.

Given the identification of *CLK1* exon 4 skipping as a recurrent splice event across the cohort, we next examined whether exon 4 inclusion (PSI) and total *CLK1* expression were associated with survival outcomes. Using multivariate Cox proportional hazards models adjusted for cluster membership, extent of resection, histology, and age at diagnosis, we found that higher *CLK1* exon 4 PSI was independently associated with improved overall survival (HR = 0.17, p < 0.01; **Figure 3F**). Tumor resection remained a favorable prognostic factor, whereas non-LGG tumors and Cluster 7 membership were associated with worse outcomes, consistent with prior analyses.

In contrast, when examining *CLK1* expression, we observed cluster-specific associations with survival. In Clusters 3 and 7—enriched for *BRAF* V600E–mutant low-grade gliomas (Cluster 3) and for other high-grade gliomas, including diffuse midline gliomas (Cluster 7) – either higher *CLK1* exon 4 containing transcript abundance or total *CLK1* transcript abundance – was associated with worse overall survival after adjusting for resection status, histology, and age (HR = 1.16 in Cluster 3 and 5.96 in Cluster 7, p < 0.01; **Figures 3G–H**). These findings indicate that exon-level splicing regulation and gene-level expression of *CLK1* have distinct and lineage-associated relationships with clinical outcome.

Consistent with these observations, analysis of spliceosome pathway activity within Cluster 7 revealed that lower GSVA scores were associated with improved overall and event-free survival, whereas higher scores were associated with poorer outcomes (**Figures 3I, S4D**). Together, these results suggest that while elevated *CLK1* expression and spliceosome pathway activity track with adverse prognosis in specific high-risk clusters, exon 4 inclusion represents a distinct splicing feature whose association with outcome is independent of overall expression levels.

To further resolve the relationships between *CLK1* exon-level splicing, gene expression, and global splicing burden, we examined correlations between *CLK1* transcript abundance (TPM), exon 4 PSI, and SBI across splicing-defined clusters and histologies (**Figure S5A-C**). *CLK1* expression and SBI showed strong, cluster-dependent relationships, with positive correlations in Clusters 7, 8, and 9 and a strong inverse correlation in Cluster 1 (**Figure S5A**). When stratified by histology, *CLK1* expression was positively correlated with SBI in ATRT, DMG, medulloblastoma, meningioma, other HGGs, and schwannoma, but negatively correlated in glioneuronal tumors (**Figure S5B**). In contrast, *CLK1* exon 4 PSI was strongly and positively correlated with *CLK1* expression across all clusters and histologies (**Figure S5C**). Together, these analyses indicate that exon 4 inclusion is tightly coupled to *CLK1* expression, whereas the relationship between *CLK1* expression and global splicing burden is context dependent and varies across tumor histologies.

The uniform coupling between *CLK1* exon 4 inclusion and expression across tumor contexts, together with the lineage-specific relationship between *CLK1* expression and splicing burden, prompted us to next examine whether exon 4 inclusion reflects a developmentally regulated splicing state.

### *CLK1* exon 4 inclusion marks an oncofetal, lineage-associated splicing state with context-specific sensitivity to CLK inhibition

To contextualize our findings within normal neurodevelopment, we examined *CLK1* exon 4 inclusion in non-tumor brain tissues using RNA-seq data from the Evolutionary Developmental (Evo-Devo) Atlas (N = 59) and GTEx (N = 2,642). *CLK1* exon 4 inclusion was significantly higher in pediatric brain tumors compared to GTEx adult brain, was comparable to fetal Evo-Devo samples, and decreased with age across both Evo-Devo and the PBTA (p < 0.05; **Figure 4A; Table S5E**). This pattern is consistent with an oncofetal splicing program^36^, and supports the model that pediatric CNS tumors re-engage developmentally-regulated *CLK1* splicing states that are normally attenuated during postnatal brain maturation. Reactivation of this state may impose distinct post-transcriptional regulatory requirements that are not uniformly buffered across lineages.

**Figure 4.**
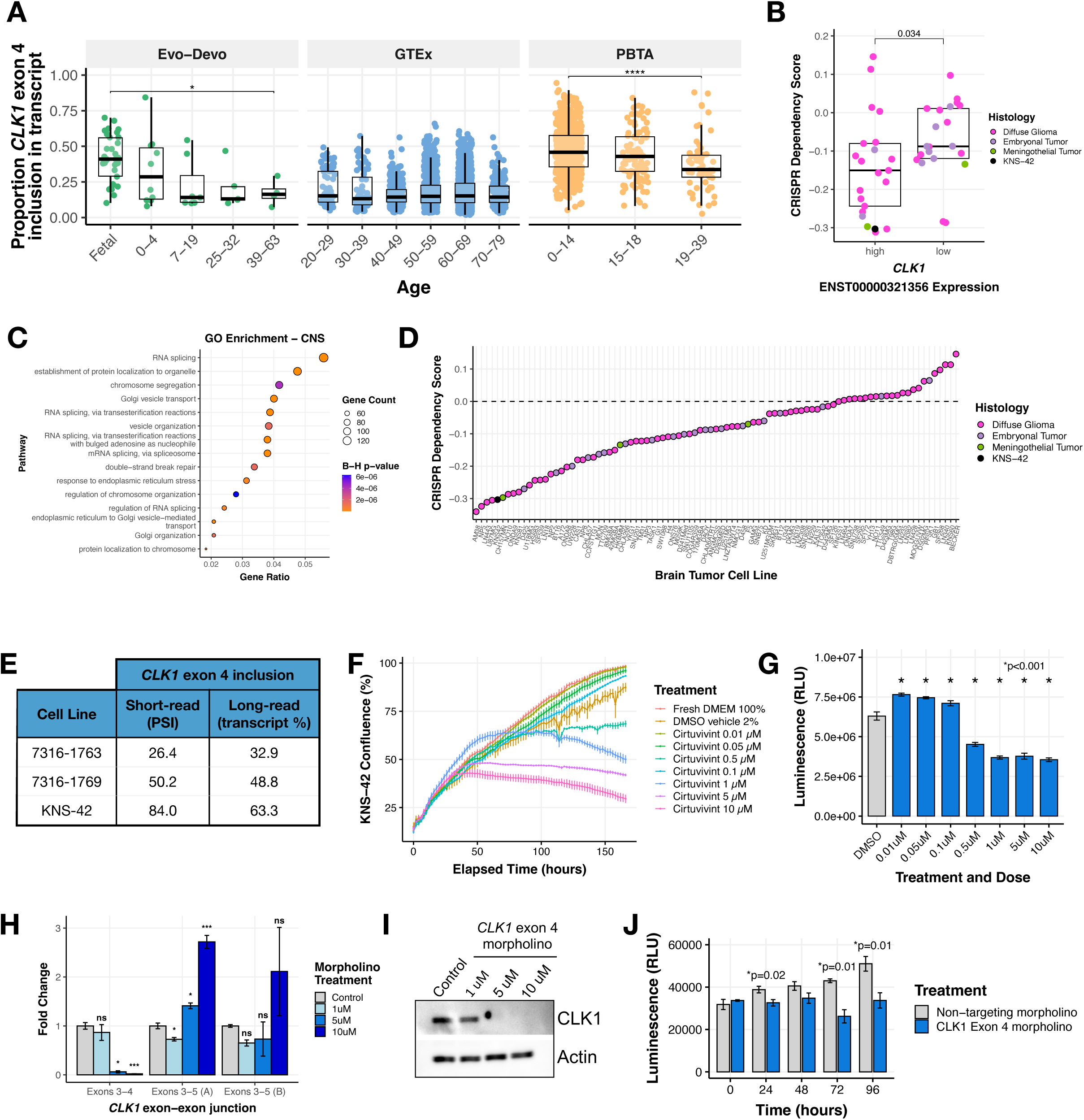
*CLK1* exon 4 is required for pediatric brain tumor cell line growth and viability. **(A)** Boxplots of CLK1 exon 4 inclusion stratified by age in non-tumor RNA-seq brain tissue data from the Evolutionary Developmental (Evo-Devo) Atlas (N = 59) and GTEx (N = 2,642). **(B)** Boxplot of DepMap dependency scores stratified by high or low *CLK1* exon 4 containing transcript expression in brain tumor cell lines. Wilcoxon p-value shown. **(C)** Dot plot of Gene Ontology (GO) gene-set enrichment analysis for genes correlated with *CLK1* across CNS cell lines from DepMap. Dot size indicates gene count size, and color represents enrichment significance. **(D)** Ranked dotplot of DepMap dependency scores in brain tumor cell lines with pediatric line KNS-42 highlighted in black. **(E)** Table showing percent spliced in for all *CLK1* exon 4 transcripts in patient-derived cell lines (7316-1763 and 7316-1769 from the CBTN) and KNS-42 (commercial) derived using short RNA-Seq or percent *CLK1* exon 4 transcripts using either long-read (ONT) sequencing. **(F)** Proliferation of KNS-42 cells treated with increasing concentrations of pan-DYRK/CLK1 inhibitor Cirtuvivint over six days. **(G)** Day 3 cell viability of KNS-42 cells treated with increasing concentrations of Cirtuvivint. Stars denote Bonferroni-adjusted p-values following pairwise Student’s *t-tests*. **(H)** Barplot showing the RNA expression fold-change in cells treated with control morpholino or morpholino targeting the *CLK1* exon 3-4 junction or exon 3-5 junction. **(I)** Western blot of CLK1 with increasing morpholino treatment of 1, 5, and 10 μM. **(J)** Cell viability of cells treated with *CLK1* exon 4 morpholino or non-targeting morpholino. Stars denote within-time paired Student’s *t-tests*.

Next, we evaluated whether *CLK1* and its exon 4 inclusion is associated with selective sensitivity in pediatric brain tumors. Analysis of the Cancer Dependency Map (DepMap) revealed that CNS tumor cell lines with high expression of the exon 4-included *CLK1* transcript ENST00000321356 (used as a proxy for exon 4-containing isoform abundance) exhibit significantly greater CRISPR sensitivity, reflected by more negative Chronos gene effect scores (0 ∼ nonessential, -1 ∼ common essentials), compared to cell lines with low exon 4 inclusion (Wilcoxon p = 0.034; **Figure 4B**). Notably, this association was restricted to CNS and myeloid malignancies and was not observed across other cancer types (**Figure S6A**), indicating lineage-specific sensitivity to perturbation of *CLK1*.

To investigate molecular features associated with this context specificity, we performed a correlation-based analysis of gene expression with the *CLK1* exon 4-containing transcript across CNS and myeloid cell lines from DepMap. We identified genes whose expression levels were significantly correlated with *CLK1* within each lineage. While only 3.9% of *CLK1-*correlated genes were shared between the two lineages (**Figure S6B, Table S4C**), both the shared core and the substantially larger lineage-specific gene sets were significantly enriched for RNA splicing, spliceosome-associated pathways, and chromatin remodeling pathways (B-H adjusted p < 0.05; **Figure S6C, 4C**). These findings suggest that *CLK1* sensitivity arises in cellular contexts characterized by coordinated, regulated exon-level splicing programs embedded within broader transcriptional and epigenetic networks, rather than simply elevated spliceosome component expression or generalized spliceosome activation.

Among pediatric CNS tumor cell lines in DepMap, the diffuse hemispheric glioma line KNS-42 exhibited the strongest sensitivity to *CLK1* perturbation (**Figure 4D**) and was therefore selected for mechanistic interrogation. KNS-42 also demonstrates robust inclusion of *CLK1* exon 4, consistent with the splicing patterns observed in our primary tumor cohort. To extend beyond a single model, we additionally evaluated two patient-derived pediatric brain tumor cell lines (7316-1763 and 7316-1769) that similarly exhibit high *CLK1* exon 4 inclusion with short-read RNA-Seq. Using Oxford Nanopore long-read RNA sequencing, we confirmed that these models express comparable ratios of *CLK1* isoforms that include or skip exon 4, validating the splice event initially identified by short-read RNA sequencing (**Figure 4E**).

Next, we tested the impact of CLK1 inhibition in KNS-42 cells using the pan-Dyrk/Clk inhibitor cirtuvivint (SM08502)^19^. Using the IncuCyte Live Cell Analysis System to monitor real-time proliferation, we observed a significant reduction in cell growth at multiple concentrations over a 6-day period (**Figure 4F**). Additionally, we observed a dose-dependent decrease in cell viability using CellTiter-Glo at three days (**Figure 4G**) and six days (**Figure S6D**) post-treatment of 0.5, 1, 5, and 10 μM Cirtuvivint. Although the reduction in proliferation/viability with cirtuvivint was dose-dependent and statistically significant, the magnitude was modest. Given that cirtuvivint is a pan-CLK/DYRK inhibitor, we next performed exon 4-specific perturbation experiments to more directly test the contribution of the catalytically active CLK1 isoform.

*CLK1* regulates the SR family of splicing factor proteins through hyper-phosphorylation of the SR-rich peptide regions of SR proteins to induce cooperative RNA binding and increased activity^33–35^. We therefore postulated that exon 4 inclusion is required to produce a stable, full-length, catalytically active CLK1 protein isoform. To directly test this hypothesis, we modulated *CLK1* exon 4 splicing using targeted morpholino oligomers (see **Online Methods**), in which we forced exon 4 skipping in the KNS-42 cell line. We performed qRT-PCR and observed a near total loss of the *CLK1* exon 4 inclusion transcript at both 5 and 10 μM of exon 4 targeted morpholino, evidenced by reduced expression of the exon 3-4 junction. At these same concentrations, we observed increased *CLK1* exon 4 skipping using primers targeting the exon 3-5 junction (**Figure 4H**). Importantly, forced *CLK1* exon 4 skipping resulted in near complete loss of CLK1 protein at 5 and 10 μM (**Figure 4I**), corroborating previous work that *CLK1* exon 4 is required for full-length and catalytically active *CLK1*^37–39^. Next, we assessed the functional impact of *CLK1* exon 4 splicing using CellTiter-Glo and confirmed that cells with high *CLK1* exon 4 skipping (*CLK1* exon 4 targeting morpholino) exhibited significantly decreased viability compared to those with *CLK1* exon 4 inclusion (non-targeting morpholino) at 24, 72, and 96 hours (p ≤ 0.01, within-time Student’s *t-test*, **Figure 4J**). Together, these data support a model in which *CLK1* exon 4 inclusion marks a lineage- and context-dependent splicing state associated with selective sensitivity to *CLK1* perturbation in the pediatric KNS-42 model tested.

To identify transcriptional and splicing changes associated with forced *CLK1* exon 4 splicing, we performed RNA-seq from KNS-42 cells treated with morpholino oligomers (N = 3 controls, N = 3 targeted to skip exon 4). We performed differential gene expression (DE) analysis and identified 1,322 genes with differential expression (569 upregulated, 753 downregulated) between cells treated with morpholino or non-targeting control (**Figure 5A, Table S6A**), including 78 oncogenes or tumor suppressor genes (TSGs, **Figure 5B**). Next, we identified a total of 4,001 unique differential splicing (DS) events (SE = 2,256; Mutually Exclusive [MXE] = 926; A5SS = 181; A3SS = 271; and RI = 367; **Table S6B-G**), including 267 oncogenes or TSGs (**Figure 5B**). There were 120 genes (2.2%) which were both DE and DS (**Figure 5D, Table S6H**), indicating these may impact total protein abundance in tumors. DS genes were significantly over-represented for mitotic spindle, E2F targets, G2M checkpoint, and nucleotide excision repair pathways (Bonf-adj p < 0.05, **Figure 5E**). To further investigate the impact on DNA repair and other pathways, we performed GSVA of DNA repair and cancer signaling pathways using DS oncogenes and TSGs and found that depletion of CLK1 was associated with upregulation of stress- and cytokine-associated signaling pathways, including TNFA/NFkB, PI3K/AKT/MTOR, IL6/JAK/STAT3, alongside downregulation of multiple DNA repair pathways, consistent with a possible metabolic and cell-state transition. (**Figures 5F-G and S5E-F**). Importantly, *CLK1* morpholino-mediated skipping resulted in significant downregulation of *CLK1* and known downstream targets, including *LRP5, AXIN2,* and *LEF1* (Log_2_FC < -0.1, B-H p-adj < 0.05, **Table S6I**), accompanied by suppression of WNT signaling (**Figure 5F-G**). While prior studies have primarily examined pharmacologic *CLK1* inhibition, the affected pathways observed here are directionally concordant with reported pathway-level effects of the pan-CLK/DYRK inhibitor cirtuvivint (SM08502)^19^, suggesting that exon 4-dependent perturbation of *CLK1* engages similar downstream regulatory programs.

**Figure 5.**
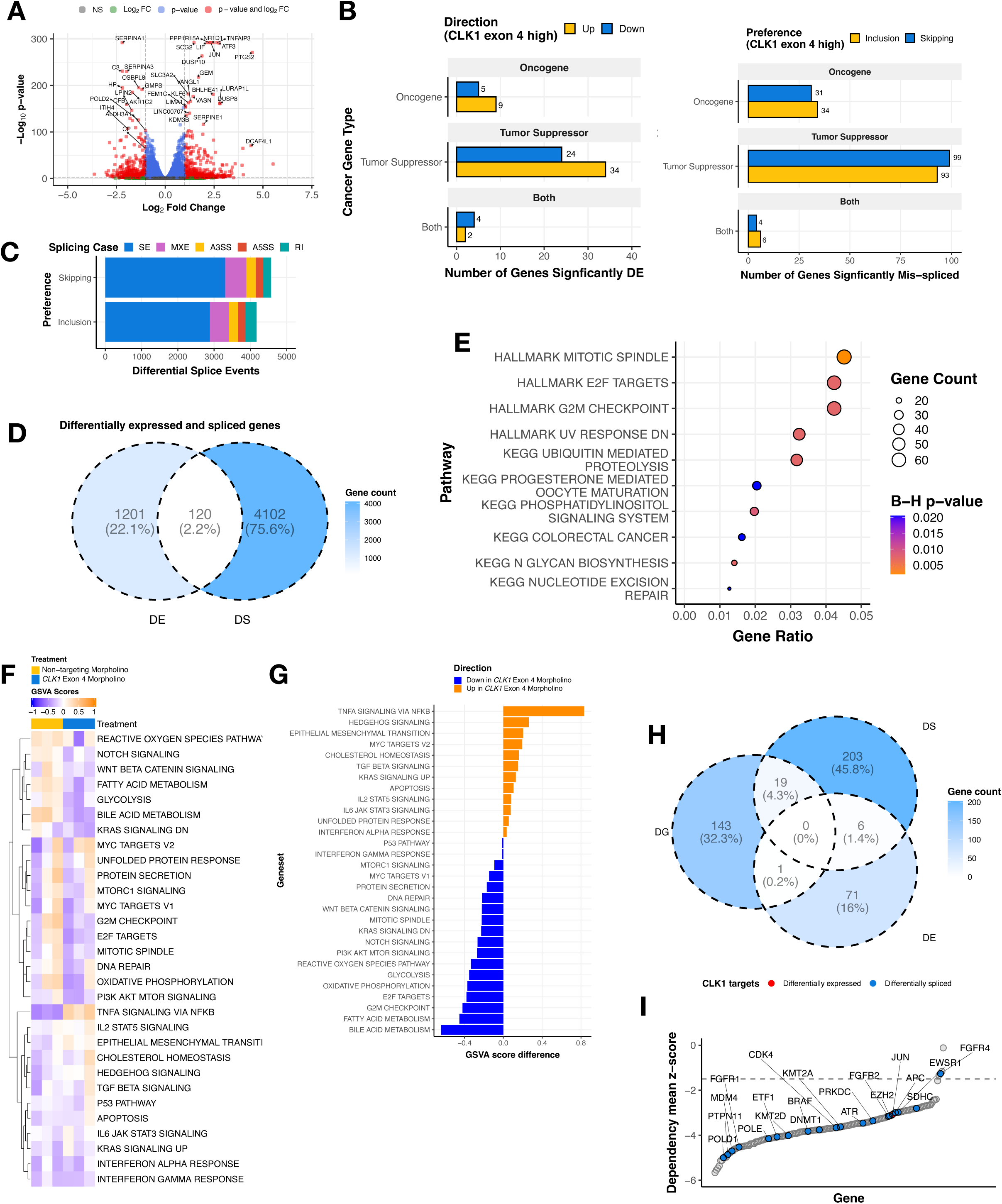
Perturbing *CLK1* disrupts RNA splicing and impairs oncogenic transcriptional programs. **(A)** Volcano plot illustrating genes differentially-expressed in KNS-42 cells treated with *CLK1* exon 4 targeting morpholino compared to cells treated with non-targeting morpholino. **(B)** Stacked barplot showing differential splicing in KNS-42 cells treated with *CLK1* exon 4 targeting morpholino compared to cells treated with non-targeting morpholino categorized by splicing type (SE, MXE, A3SS, A5SS, and RI). **(C)** Barplots displaying number of differentially expressed (DE) genes or differentially spliced (DS) genes affecting functional sites categorized by gene family.**(D)** Venn diagram depicting overlap of all DS and DE genes **(E)** Over-representation analysis of HALLMARK and KEGG pathways for DS cancer genes. Dot size indicates gene count size, and color represents enrichment significance. **(F)** Heatmap displaying single-sample HALLMARK GSVA scores for DS genes affecting functional sites in cells treated with *CLK1* exon 4 morpholino or non-targeting morpholino. **(G)** Barplots illustrate the mean GSVA score difference by treatment (n = 3 replicates per treatment). **(H)** Venn diagram depicting overlap of DS and DE genes and significant (Wald FDR < 0.05, z-score < -1.5) dependency genes (DG) identified in matched CBTN HGG cell lines through CRISPR dependency experiments from the Childhood Cancer Model Atlas (CCMA v3). **(I)** Ranked dotplot of significant CRISPR gene dependency mean z-scores for pediatric HGG cell lines with *CLK1* expression and splicing-based target genes highlighted in red and blue, respectively.

Finally, we asked whether *CLK1 exon 4* inclusion levels are associated with changes in essential oncogenes defined by the pediatric gene dependency maps of the Childhood Cancer Model Atlas^40^. We identified 20 such genes that also exhibit significant gene dependencies in PBTA cell lines (**Figure 5H-I, Table S6J**), including seven regulators of MAPK signaling: *BRAF, EZH2, RAF1, JUN, FGFR1, FGFR2,* and *SRC*. For instance, expression of mRNAs encoding proto-oncogene *SRC*^41,42^ was higher in cells with high *CLK1* exon 4 inclusion (non-targeting morpholino), consistent with a role for *CLK1*-associated splicing states in modulating *SRC* expression. Differential splicing effects were more complex, as they involved multiple transcripts within individual genes; however, taken together, these data suggest that transcript-level changes associated with differential *CLK1* exon 4 inclusion may contribute to sensitivities in pathways such as DNA repair and MAPK signaling. This interpretation is consistent with established links between alternative splicing with cancer progression^43–45^.

While extending functional validation to additional pediatric brain tumor models, non-malignant CNS counterparts, and *in vivo* systems will be critical for defining therapeutic windows and translational relevance, these findings position *CLK1* exon 4 as a candidate lineage-associated axis of splicing regulation with potential clinical relevance in pediatric CNS tumors.

## Discussion

Pediatric brain tumors remain the leading cause of disease-related mortality in children. In this study, we perform a large-scale, cross-histology analysis of alternative splicing across pediatric CNS tumors and demonstrate that splicing dysregulation is pervasive, heterogeneous, and clinically informative. By integrating exon-level splicing, gene expression, pathway activity, proteogenomics, and functional perturbation, our work provides a framework for uncovering how splicing programs contribute to tumor biology beyond canonical genomic alterations.

We introduce the Splicing Burden Index (SBI) as a quantitative, sample-level metric that enables comparison of differential splicing across tumors in a cohort without reliance on matched normal tissue, an important consideration when normal brain tissue is difficult to obtain. SBI varied across and within tumor histologies, highlighting substantial inter-tumoral heterogeneity in splicing programs. Contrary to our initial hypothesis, tumor mutational burden (TMB) showed only a very weak association with SBI, explaining little variance and arguing against splicing dysregulation serving primarily as a compensatory mechanism in genetically quiet tumors. Instead, global splicing burden appears to reflect context-dependent dysregulation of RNA processing programs.

Clustering tumors by highly variable splice events identified ten splicing-defined groups enriched for specific histologies and molecular subtypes. Several clusters remained independently associated with survival after adjustment for histology and clinical covariates, demonstrating that splicing captures prognostically-relevant heterogeneity beyond historical tumor classification alone. Importantly, the relationship between splicing burden and outcome was context dependent, with cluster-specific interactions indicating that the biological consequences of splicing dysregulation are not uniform across pediatric CNS tumors.

Pathway-level analyses further underscored this complexity. Spliceosome pathway activity varied across splicing-defined clusters and was independently associated with adverse outcomes in multivariate models. However, spliceosome GSVA scores were not correlated with SBI, indicating that pathway activation and splicing burden represent distinct dimensions of splicing dysregulation. Integration with matched proteogenomic data confirmed that transcript-based GSVA scores reflect underlying protein abundance, supporting their biological relevance.

Mechanistically, deleterious coding mutations in spliceosome components and splicing factor genes were rare, whereas expression changes in splicing regulators were widespread, including across SRSF and hnRNP family members. These findings support a model in which altered expression of splicing regulators, rather than recurrent genetic disruption of splicing machinery, contributes to widespread splicing alterations in pediatric CNS tumors.

To identify splice events with potential functional relevance, we prioritized recurrent differential splicing events predicted to alter annotated protein features. This approach highlighted *CDC-like kinase 1 (CLK1)* as a compelling candidate, based on a recurrent event affecting exon 4, which is required for canonical kinase activity. Although most tumors exhibited high exon 4 inclusion, a subset showed significant skipping, motivating further investigation. Integration with developmental transcriptomes further demonstrated that *CLK1* exon 4 inclusion follows an oncofetal pattern, supporting a model in which pediatric CNS tumors re-engage developmentally regulated splicing states that may create context-specific sensitivities.

Clinical analyses revealed that exon-level splicing and gene-level expression of *CLK1* have distinct and context-dependent associations with outcome. Higher exon 4 inclusion was independently associated with improved survival across the cohort, whereas elevated *CLK1* expression tracked with worse outcomes in specific high-risk clusters. Correlation analyses clarified these relationships: exon 4 inclusion was tightly coupled to *CLK1* expression across all contexts, while the relationship between *CLK1* expression and global splicing burden varied by cluster and histology. These findings highlight the importance of distinguishing isoform-level regulation from bulk expression and pathway activity.

Functional perturbation of *CLK1* exon 4 in a pediatric HGG model reduced overall *CLK1* protein abundance, impaired cell viability, and induced widespread transcriptional and splicing changes affecting cell cycle, DNA repair, and MAPK signaling pathways. While limited to a single pediatric tumor model, these experiments provide proof-of-concept evidence that exon-level disruption of *CLK1* can perturb oncogenic programs. The modest phenotype observed with pan-CLK/DYRK inhibition highlights both the challenges of polypharmacologic targeting and the importance of isoform-resolved perturbation strategies when evaluating splicing-associated functional sensitivities.

Notably, *CLK* family kinases, including *CLK1*, are already under clinical investigation in multiple adult malignancies through early phase trials. The Pan-Clk/Dyrk Inhibitor cirtuvivint (SM08502) is being used in a phase 1 clinical trial in patients with acute myeloid leukemia (AML) and myelodysplastic syndromes (MDS)^46^, and has shown preclinical efficacy in non-CNS solid tumors such as triple negative breast cancer, pancreatic cancer, castrate-resistant prostate cancer, colorectal cancer, endometrial cancer, and non-small cell lung cancer^14,15,16–20^. An ATP-competitive, macrocyclic inhibitor of the CLK family, BH-30236^47^, is in a Phase 1/1b clinical trial for patients with AML and MDS^48^. Finally, the CLK1-specific inhibitor, CTX-712, is in Phase 1/2 trial for relapsed or refractory AML and high risk MDS^49^, underscoring *CLK1* and its family as a target across diverse tumor histologies. Our study extends the rationale for CLK1 inhibition to pediatric brain tumors, particularly those with high splicing burden and exon 4 inclusion.

In summary, our work provides a systematic, integrative view of alternative splicing across pediatric CNS tumors, identifies splicing-informed tumor clusters with clinical relevance, and prioritizes *CLK1* exon 4 inclusion as a recurrent, developmentally-regulated splice event candidate for further preclinical testing. By distinguishing exon-level regulation from global splicing burden and pathway activation, this study highlights the importance of isoform-resolved analyses and provides a foundation for future splicing-directed therapeutic strategies in pediatric brain cancer.

### Limitations of the study

In this study, initial splicing quantifications relied on short-read RNA-Seq technology, which limits resolution of full-length isoforms and complex multi-exon events. Future integration with long-read RNA-Seq will enable more comprehensive isoform-level analyses^50^. The absence of matched normal RNA for each tumor of origin tissue restricts the ability to define mutually exclusive or tissue-specific splicing events. For example, within histologies (eg: LGG), the primary site of the tumor can vary widely depending on diagnosis and it would be ideal to match each tumor to its tissue of origin. We mitigated these limitations by leveraging multiple non-tumor reference cohorts, developmental datasets, and the SBI metric. Finally, while functional validation supports a role for *CLK1* exon 4 in tumor cell fitness, extending these studies to additional pediatric models, non-malignant CNS counterparts, and *in vivo* systems will be essential to define therapeutic windows and translational relevance.

## Author Contributions

Conceptualization: ASN, JLR, ACR, RJC, PJS

Methodology: ASN, JLR, RJC, ACR, KSR, AK, SA, KLC, PJS

Software: BE, ASN, RJC, PJS, JLR

Validation: PS, KC, RJC, KLC, KEH, ASN, AL, PS, PJS

Formal Analysis: ASN, JLR, RJC, KLC, KSR, BE, PJS, AS

Investigation: ASN, JLR, RJC, KSR, AK, PJS

Resources: JLR, ACR, ATT, PBS, CB, SJD, Kal LC, DH

Data Curation: JLR, ASN, RJC

Writing - Original Draft: ASN, JLR, KSR, PS, KLC, PJS

Writing - Review & Editing: ASN, JLR, RJC, SJD, KLC, JBF, JMD, PJS

Visualization: ASN, JLR, KSR, RJC, KLC, PJS

Supervision: JLR, ACR, ATT

Project Administration: JLR, ACR

Funding Acquisition: JLR, ACR, PBS, ATT, SJD

## Ethics statement

This study utilized publicly available genomic data from the Open Pediatric Cancer (OpenPedCan) project, derived from the Children’s Brain Tumor Network (CBTN) and the Pediatric Neuro-Oncology Consortium (PNOC). These consortia have established protocols for the ethical collection and sharing of patient data. The patient-derived cell lines 7316-1763 and 7316-1769 were obtained via request from the CBTN, and the KNS-42 cell line was purchased commercially. While this study did not directly involve the collection of new human or animal subject data, the use of these pre-existing resources implies that the original data collection was conducted with appropriate institutional review board approval and informed consent procedures.

## Data Availability

### Patient genomic data

All pediatric brain tumor raw data are available upon request from the database of Genotypes and Phenotypes (dbGAP), accession number phs002517.v2.p2, and/or from the Children’s Brain Tumor Network (https://cbtn.org) and the Pacific Pediatric Neuro-Oncology Consortium (pnoc.us) for data not immediately available in dbGaP. All processed data used in this study were derived from the OpenPedCan project^21^ v13 data release at https://github.com/d3b-center/OpenPedCan-analysis. All code for the manuscript analyses and figures are openly available at https://github.com/rokitalab/clk1-splicing.

### Non-tumor control RNA-Seq

RNA-Seq data from the GTEx project (dbGAP Accession phs000424) and the Evo-Devo atlas (Array Express Accession E-MTAB-6814) were harmonized with GENCODE v39 using the Kids First Data Resource Center workflow at https://github.com/kids-first/kf-rnaseq-workflow.

### CLK1 morpholino RNA-Seq

RNA-sequencing data from the *CLK1* morpholino experiment has been deposited in GSE273841.

### Merged primary and summary data

Merged primary matrices and summary files utilized in this manuscript were derived from are openly accessible via the download script in the https://github.com/rokitalab/clk1-splicingrepository.

## Supporting information

SuppInfo

SuppFigs

## Acknowledgements

We graciously thank all of the patients and families of the Children’s Brain Tumor Network (CBTN) and the Pediatric Neuro-oncology Consortium for donating tissue which enabled this research. The authors thank the following collaborators who contributed experiments, analyses, code, and/or code review that were ultimately not included in the manuscript: Poonam Sonawane, Cullen Wilson, Krutika S. Gaonkar, and Run Jin.

