## Supplementary material for "Alternative splicing in pediatric central nervous system tumors highlights oncofetal candidate *CLK1* exon 4": SuppInfo

### Supplemental Figure Legends

**Figure S1. (A)** UpsetR plot of recurrent differential splicing events (N ≥ 2 of samples within a histology). Correlation plots for SBI vs TMB across the entire cohort including **(B)** and excluding **(C)** hypermutant and ultra-hyper mutant tumors. **(D)** Correlation plots separated by histology. All plots show Pearson's R and p-values.

**Figure S2. (A)** Enrichment heatmaps for all histologies, **(B)** MB subtypes, **(C)** HGG subtypes by cluster. Boxes contain N tumors, Odds ratio (OR) in parentheses, and stars denote Fisher's Exact test FDR < 0.05. **(D)** Stacked barplot showing tumor histology membership in each cluster stratified by molecular subtype for EPN, HGG, LGG, and MB histologies. **(E)** Boxplots of splicing burden index by cluster, colored by histology. All boxplots represent the 25th and 75th percentiles, and the bar represents the median.

**Figure S3. (A)** Kaplan-Meier plot for OS by cluster membership **(B)** Forest plot of cox proportional hazards multivariate OS model of cluster membership with covariates tumor resection, KEGG spliceosome GSVA score quartile, and age at diagnosis. Black and white diamonds indicate statistically significant and not significant HRs, respectively, with intervals denoting 95% confidence intervals. Gray diamonds indicate reference levels of factor covariates. **(E)** Correlation plot of spliceosome GSVA scores and matched mean proteomic score of associated proteins, colored by histology. **(F)** Correlation plot of SBI and spliceosome GSVA scores.

**Figure S4. (A)** Oncoprint displaying mutation frequencies of recurrent canonical cancer driver or SF genes in samples with annotations for cluster membership, gender, cancer predisposition, histology, CNS region, and tumor mutation status. **(B)** Boxplots of *PRKDC* exon 80 inclusion stratified by age in non-tumor RNA-seq brain tissue data from the Evolutionary Developmental (Evo-Devo) Atlas (N = 59) and GTEx (N = 2,642), and tumor samples from PBTA polyA (N = 557) and unstranded cohorts (N = 719). **(C)** Boxplots of *CLK1* exon 4 PSI values by histology. **(D)** Kaplan–Meier curves for overall survival (EFS) in Cluster 7, stratified by GSVA score.

**Figure S5.** Scatter plots showing correlations between *CLK1* exon 4 containing transcript abundance (TPM) and splicing burden index (SBI) across **(A)** splicing-defined clusters colored by histologies and **(B)** tumor histologies.

**(C)** Scatter plots showing correlations between exon 4 PSI and *CLK1* exon 4 containing transcript abundance (TPM) across clusters, colored by histologies.

**Figure S6. (A)** Boxplot of DepMap dependency scores stratified by high or low *CLK1* exon 4 containing transcript expression across all available DepMap tumor cell lines. Wilcoxon p-value shown. **(B)** Venn diagram of the overlap of *CLK1*-correlated transcripts across CNS and myeloid malignancies from DepMap. **(C)** Dot plot of Gene Ontology (GO) gene-set enrichment analysis for genes correlated with *CLK1* shared between CNS and myeloid cell lines from DepMap. Dot size indicates gene count size, and color represents enrichment significance. **(D)** Cell viability assay after six days of treatment of KNS-42 cells with increasing concentrations of pan-DYRK/CLK1 inhibitor Cirtuvivint. **(E)** Heatmap presenting single-sample DNA repair pathway GSVA scores for DS genes affecting functional sites in cells treated with *CLK1* exon 4 morpholino or non-targeting morpholino. **(F)** Barplots displaying mean DNA repair pathway GSVA scores (n = 3 replicates per treatment).

### Supplementary Table Legends

**Table S1**: Sample metadata. **(A)** Readme and feature definitions. **(B)** Sample information with clinical metadata and demographics **(C)** CNS region definition from OpenPedCan[1].

**Table S2**: Histology-specific splicing events. **(A)** Differential splicing events. SpliceID includes gene name and event coordinates as provided by rMATs. **(B)** Cluster membership for each sample. **(C)** All nominally ($p < 0.05$) and significantly ($p$ adj $< 0.05$) differentially-expressed pathways by cluster.

**Table S3**: **(A)** KEGG spliceosome gene list. **(B)** HUGO spliceosome component gene list **(C)** Splicing factor and related genes from Sebestyén, et. al[2]. **(D)** Somatic mutations in HUGO spliceosome genes or **(E)** splicing factor and related genes across the cohort. **(F)** DeSeq2 results comparing high- vs low-SBI tumors for splicing factors.

**Table S4:** Pearson correlation coefficients between splicing factor gene (**Table S3C**) transcripts per million (TPM) and SBI by **(A)** cluster, and **(B)** histology. **(C)** Correlation between *CLK1* ENST00000321356 and all other transcripts inDepMap cell lines with CNS or myeloid malignancies.

**Table S5**: Differential splicing events impacting functional sites. **(A)** Skipping (dPSI < -2 z-score) DS events. **(B)** Inclusion (dPSI > 2 z-score DS events. **(C)** Functional splice variants subsetted for known kinases that are also splicing factors[2]. **(D)** *CLK1* exon 4 PSI for each sample in the cohort. **(E)** Wilcoxon rank-sum test results for age-stratified comparisons of *CLK1* exon 4 inclusion among the study cohort, GTEx, and Evo–Devo normal samples. P-values are Benjamini–Hochberg-adjusted.

**Table S6**: *CLK1* morpholino analyses. **(A)** Differential gene expression results from DeSeq2 and **(B)** rMATs results comparing treated with *CLK1* exon 4 morpholino and non-targeting morpholino. **(C)** Differential splicing events associated with SE that correspond to known Uniprot functional sites. **(D)** Differential splicing events associated with A5SS that correspond to known Uniprot functional sites. **(E)** Differential splicing events associated with A3SS that correspond to known Uniprot functional sites. **(F)** Differential splicing events associated with RI that correspond to known Uniprot functional sites. **(G)** Differential splicing events associated with MXE that correspond to known Uniprot functional sites. **(H)** Genes that are differentially expressed and spliced. **(I)** Differentially expressed *CLK1* target genes overlapping with known *CLK1* targets. **(J)** Differentially expressed or differentially spliced *CLK1* target genes overlapping essential oncogenes defined by CCMA v3. **(K)** *CLK1* exon 3-4, exon 3-5, and exon 3-5 junction forward and reverse primer sets.

### <u>Online Methods</u>

#### Molecular tumor classification

Tumor classification and molecular subtyping for this study were derived from the v13 release of the Open Pediatric Cancer (OpenPedCan) project[1], an open, harmonized multi-omic resource that builds upon and extends the original Open Pediatric Brain Tumor Atlas (OpenPBTA)[3]. OpenPedCan integrates clinical pathology,

DNA methylation profiling, and molecular features from sequencing data to assign high-confidence tumor diagnoses and molecular subtypes.

DNA methylation profiling was used as a key component of tumor classification where available, using the v12b6 version of the DKFZ classifier[4]. However, we note that methylation-based classification may fail or yield low confidence scores in cases of low tumor purity, degraded material, or ambiguous profiles. To minimize misclassification, molecular subtypes in OpenPedCan were assigned only when supported by high-confidence methylation classification and/or concordant molecular alterations, in conjunction with the original pathological diagnosis.

Across the OpenPedCan v13 release used in this study, 655 of 729 tumors (89.8%) had matched DNA methylation data. Of these, 457 tumors (69.8%) achieved a classifier subclass score ≥ 0.8 and were not assigned to a control or ambiguous class, and were therefore considered informative for molecular subtyping. For tumors with discrepant methylation results, discordant molecular features, or inconsistencies with pathology reports, cases were reviewed with expert neuropathology input. In some instances, this process resulted in updated integrated diagnoses (e.g., reclassification from generalized high-grade glioma to diffuse midline glioma, H3 K28-altered, or from medulloblastoma NOS to molecularly defined subtypes), or complete removal of samples from downstream analyses and the Kids First Data Resource Portal due to sample misidentification.

For all tumors, the original clinical pathology diagnosis (“pathology_diagnosis”) is retained, while derived classifications such as “cancer_group” or “plot_group” are used for downstream analyses, including those presented in this study. Molecular subtypes were assigned conservatively to prioritize specificity over completeness, ensuring that subtype-based analyses reflect biologically and clinically meaningful groupings. OpenPedCan is an actively maintained and versioned resource; all analyses in this study were performed using the v13 data release to ensure reproducibility. Additional details regarding data generation, quality control, methylation classification, and integrated diagnostic review are described in the OpenPedCan project manuscript[1].

#### ***CLK1* exon 4 related visualizations and correlations**

To visualize the *CLK1* exon 4 splice event, we utilized the R package *ggsashimi*[5]. We correlated *CLK1* exon 4 PSI values with *CLK1-201* or *CLK1* TPM. We computed Pearson correlation coefficients and p-values of this plot using the R package *ggpubr*[6]. High *CLK1* exon 4 inclusion tumors were defined as those with PSI values above the 75th percentile, while low SBI samples were those with PSI values below the 25th percentile comparing across all samples.

**Splicing burden index (SBI) calculation**

The following describes the SBI calculation used in the manuscript.

Let X be the list of all samples, where $X_{ij}$ represents the j-th item in the i-th sample.

Let n be the number of items in each sample.

Let SE be the splice event of interest.

Let $SE_i$ be the number of splice events in the i-th sample.

Let $mean_{SE}$ be the mean of the splice event across all samples.

Let $\sigma_{SE}$ be the standard deviation of the splice event across all samples.

Let SBI be the proportion of splice events that have z-scores > |2| out of the total number of splice events in a particular sample.

Then the equation for SBI is:

$$SBI_i = \frac{\sum_{j=1}^{n} I\left(\left|\frac{X_{i,j} - mean_{SE}}{\sigma_{SE}}\right| > |2|\right)}{n \cdot len(X)},$$

*where* i = 1 to len(X) and j = 1 to n

Let **X** be the set of all samples.

Let $\mathbf{X_{ij}}$ represent the PSI value of the **j-th splice event (SE)** in the **i-th sample**.

Let **n** be the number of splice events measured in each sample.

Let $\mathbf{SE_i}$ be the number of observed splice events (non-missing) in sample **i**.

Let $\mathbf{meanSE_j}$ be the **mean PSI** of splice event **j** across all samples.

Let $\mathbf{\sigma SE_j}$ be the **standard deviation** of splice event **j** across all samples.

For each PSI value $\mathbf{X_{ij}}$, compute the z-score:

$$\mathbf{z_{ij} = (X_{ij} - meanSE_j) / \sigma SE_j}$$

$$\mathbf{SBI_i = [Number\ of\ splice\ events\ with\ |z_{ij}| > 2] / SE_i}$$

Or, using summation notation: $\mathbf{SBI_i = (\Sigma_{(j=1)}^{n} I(|z_{ij}| > 2)) / SE_i}$

Where:

- **I(condition)** = 1 if condition is true, 0 otherwise.
- $\mathbf{SE_i}$ = total number of valid (non-NA) splice events in sample *i*.

We compared PSI values of each primary tumor against all other tumors in the cohort. We first computed mean and standard deviation metrics for each alternative splicing event observed in at least one sample. Then for each sample in each group or histology, we identified the proportion of genes that underwent aberrant splicing as defined by a z-score > |2| across the entire transcriptome that undergoes alternative splicing.

**Consensus clustering**

We filtered the PSI matrix for splice events reported in $\geq$ 25% of samples, and performed hierarchical clustering of samples based on the top 5,000 splice events with the highest PSI variance using the Euclidean distance measure and the Ward D2 agglomeration method. We determined the optimal number of clusters using the "elbow method"; briefly, we plotted the within-cluster sum of squares (WCSS) against the cluster number, and the x-axis value at which an inflection point was observed was defined as the optimal cluster number. We observed distinct clustering of samples by RNA library type within tumor histologies, and therefore re-ran clustering analyses separately for stranded and poly-A stranded libraries. We assessed enrichment of tumor histologies and molecular subtypes within identified clusters using Fisher's exact tests, and defined significant enrichment of diagnoses within clusters at FDR < 0.05.

**Clustering-based differential expression or pathway enrichment**

We performed gene set variation analysis (GSVA) on expression data derived from stranded RNA-seq libraries to calculate GSVA scores for KEGG spliceosome and HALLMARK cancer pathways using the *GSVA* R package[7]. We identified differentially expressed pathways between clusters using the *limma* R package[8], and visualized cluster differential expression using the *pheatmap* R package[9].

**Differential expression and visualization**

Differential expression was performed based on a model using the negative binomial distribution, a method employed by the R package *DeSeq2*[10]. Those differential genes that had a p-value < 0.05 were deemed as significantly up or down-regulated. Volcano plots were generated by the *EnhancedVolcano* R package. Bar plots were generated using the R package *ggplot2*[11]. Note: differential expression analyses were limited to stranded-only RNA-seq samples in order to limit batch effects.

**Identification of recurrent functional differential splicing variants in pediatric HGGs**

To identify differential or aberrant alternative splicing events, we assessed the percent spliced in (PSI) value of each splice event relative to the median PSI value of splice event across all samples. Splicing events with a ΔPSI exceeding |2| z-scores from the median PSI value were classified as differential or aberrant. For these events, we computed average ΔPSIs and generated bed files for each mis-spliced exon event. We then obtained bed files of known functional annotations as defined by Uniprot release 2025_03[12]. We ran bedtools v2.30[13] to find the overlap between mis-spliced exons and functional features using the command `bedtools intersect -wo -a`. We then plotted summary data by functional category (disulfide bonding sites, localization signals, amino acid modifications, and other).

**Upset R and Volcano plots**

To visualize the intersections of multiple sets, we employed the UpSetR[14] plot in R. The input data consisted of differential and recurrent splicing events, if it was > 2 z-scores from the meanPSI and 2% of the histology-specific cohort. Volcano plots were generated by the *EnhancedVolcano* R package.

**Splicing burden index and tumor mutation burden correlations**

We identified samples with available data for both SBI (RNA-Seq) and WGS or WXS tumor mutation burden (TMB) from OpenPedCan. Using the R package ggscatter, we performed a Pearson correlation analysis to examine the relationship between SBI and TMB. To ensure robustness, we repeated this analysis after excluding hyper-mutated samples (defined as those with TMB ≥ 10). Subsequently, we compared the distribution of TMB between high SBI and low SBI tumor samples using the Wilcoxon rank-sum test. High SBI samples were defined as those with SBI values above the 75th percentile, while low SBI samples were those with SBI values below the 25th percentile. The analyses were conducted across all samples and further stratified according to `plot_group`, as specified in the histologies clinical file.

**Pathway over-representation analysis (ORA) and gene set variation analysis (GSVA)**

We conducted over-representation analysis (ORA) using the R package clusterProfiler[15] and pathway data from the msigdbr package[16], including “CP:KEGG", "CP:BIOCARTA", “CP:HALLMARK”, and “TFT:GTRD.” After inputting the genes of interest (e.g. differentially spliced), we applied a p-value cutoff of 0.05 and used the Benjamini-Hochberg (BH) method for p-value adjustment. For visualization of the over-represented pathways, we employed the `enrichplot::dotplot()` function, displaying the gene ratio and the count of genes in each pathway.

To perform Gene set variation analysis (GSVA) we utilized the R packages `GSVA` and `msigdbr`. Expression data for our samples, sourced from OpenPedCan v13[1], were used to compute gene-set enrichment scores. Genes with zero variance were excluded from the analysis. We then assessed enrichment in Hallmark, KEGG, and custom pathways from Knijnenburg et al[17]. Gaussian-distributed scores were calculated using *gsvaParam* function in R. The results were visualized using heatmaps of GSVA scores, generated with the R packages *ComplexHeatmap* and *circlize*.

**Oxford Nanopore Technologies (ONT) Targeted Long-Read RNA-Sequencing**

We designed primers (**Table S6K**) to bind all isoforms of *CLK1* to ensure full coverage of all alternative splicing events. 5 ng of cDNA were amplified with LongAmp Taq 2X Master Mix (M0287S, New England Biolabs) for 25 cycles. The resulting amplicons were subjected to amplicon-seq (SQKNBD112.24, ONT) library preparation, loaded into a Spot-ON flow cell R9 Version (FLO-MIN112, ONT), and sequenced in a MinION Mk1C device (ONT) until at least 1,000 reads per sample were obtained.

The alignment pipeline began by processing raw Nanopore electrical signal data (FAST5 files) using the Guppy basecaller (v6.0.6) to generate sequence data (FASTQ format). These reads were then aligned to the human reference genome (GRCh38/hg38) using Minimap2 version 2.24-r1122 with the splice-aware long-read preset. The resulting aligned files underwent Quality Control (QC) using Samtools (v1.12), filtering reads based on length (≥ 200 bp) and mapping quality to ensure high-confidence input. The final alignments were then inspected visually using IGV version 2.12.3 to validate complex splicing events.

To quantify exon 4-specific splicing, reads were classified as exon 4-included or exon 4-skipped based on annotated transcripts. We computed the proportion of exon 4 inclusion transcripts relative to total *CLK1* transcripts. This computational quantification provided the values reported in **Figure 4E**.

**DepMap and CRISPR dependency analyses**

Datasets comprising gene transcript expression, cell line information, and CRISPR dependency scores were downloaded from DepMap (version 24Q2). The expression of *CLK1* ENST00000321356 (exon 4 containing transcript) was categorized into high and low TPM expression, defined by values above the 75th quantile and below the 25th quantile, respectively. CRISPR dependency scores were plotted on the y-axis, and Wilcoxon tests were conducted to compare high versus low TPM expression groups. These were stratified for each cell line type. Additionally, CRISPR dependency scores for all CNS/brain cell lines were plotted, with KNS-42 highlighted in red. For the Childhood Cancer Model Atlas CRISPR dependency analyses, we acquired data from the Childhood Cancer Model Atlas[18]. We plotted CRISPR dependency scores (z) on the y-axis for each gene in CBTN pediatric HGG cell lines, either as median scores or stratified by individual patients with genes of interest highlighted.

**Proteogenomic analysis**

Pediatric proteomics, phosphoproteomics, and RNA data were obtained from the Clinical Proteomic Tumor Analysis Consortium (CPTAC) via the ProTrack: Pediatric Brain Tumor open-source web portal. Data and z-scores were computed using the methods described by Petralia et al[19].
