## Supplementary figures and images for "Alternative splicing in pediatric central nervous system tumors highlights oncofetal candidate *CLK1* exon 4"

### SuppFigs

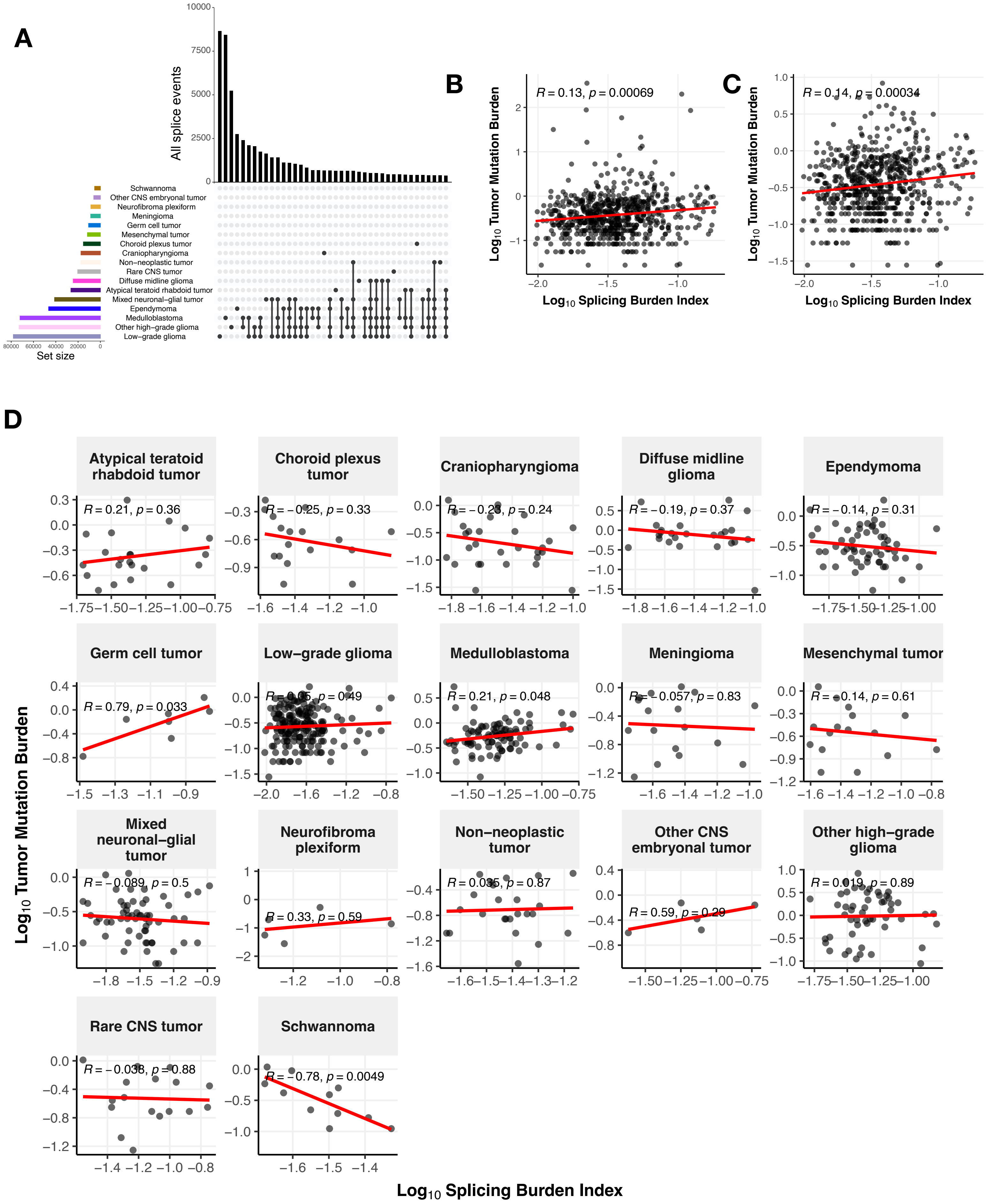

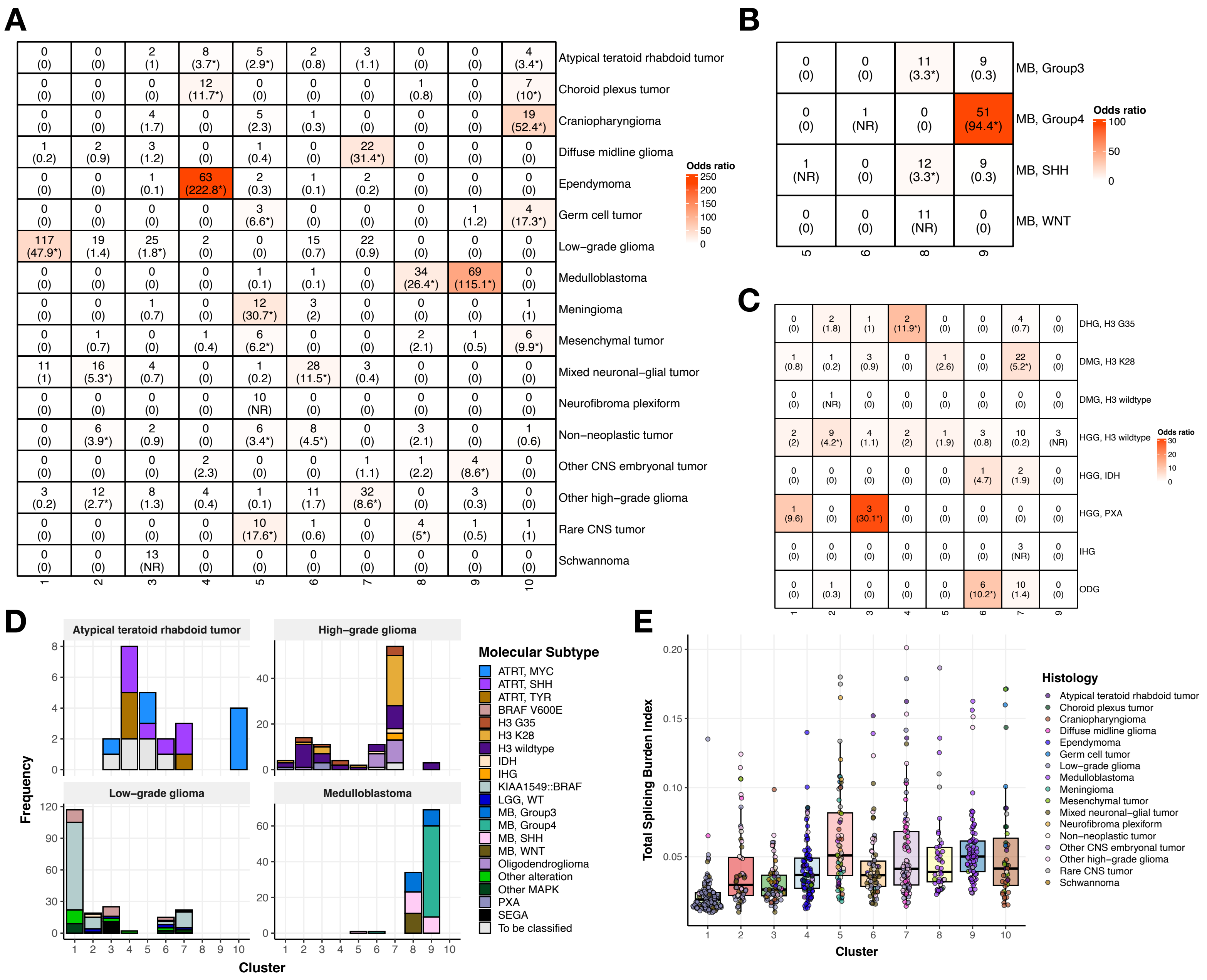

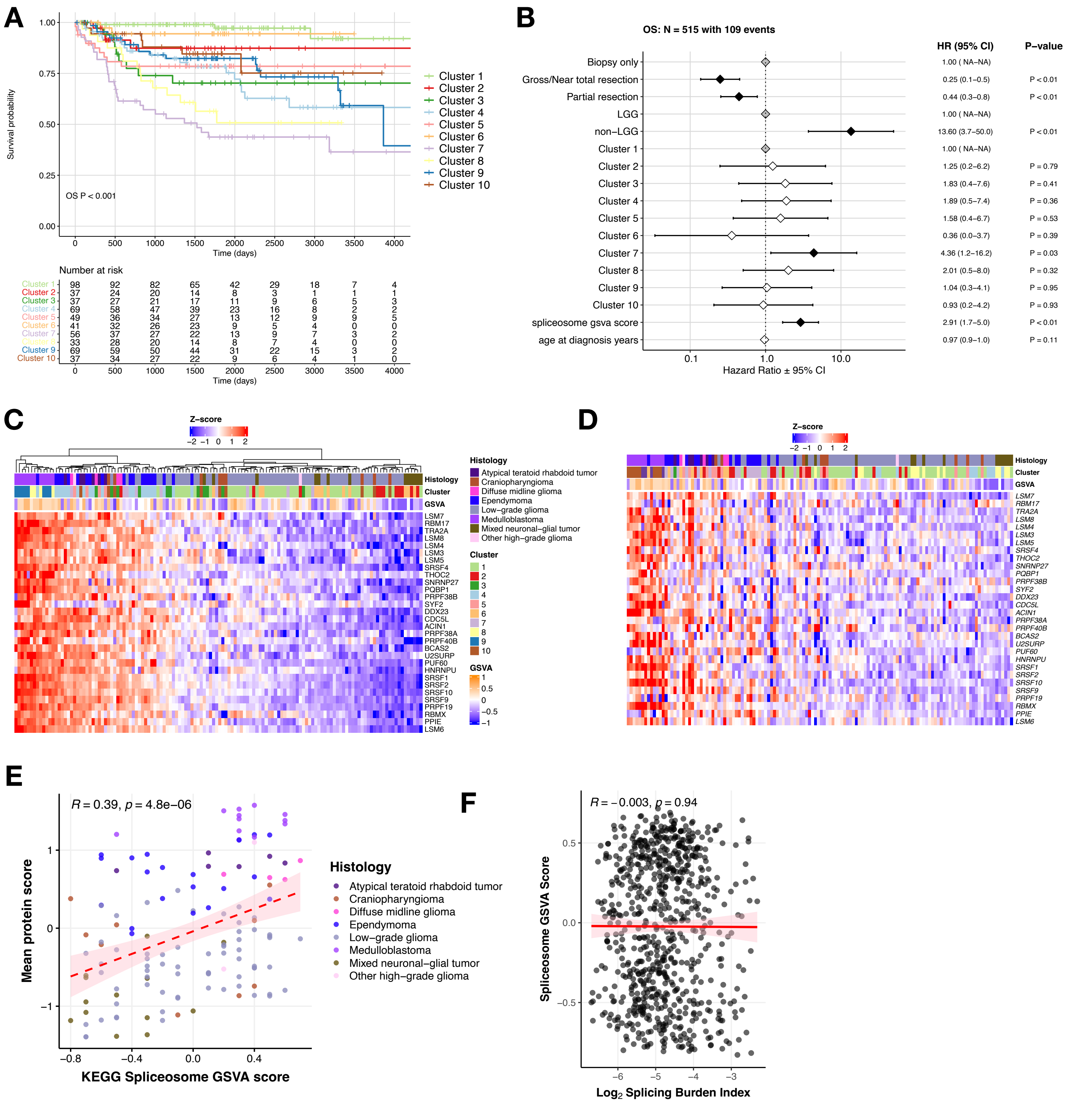

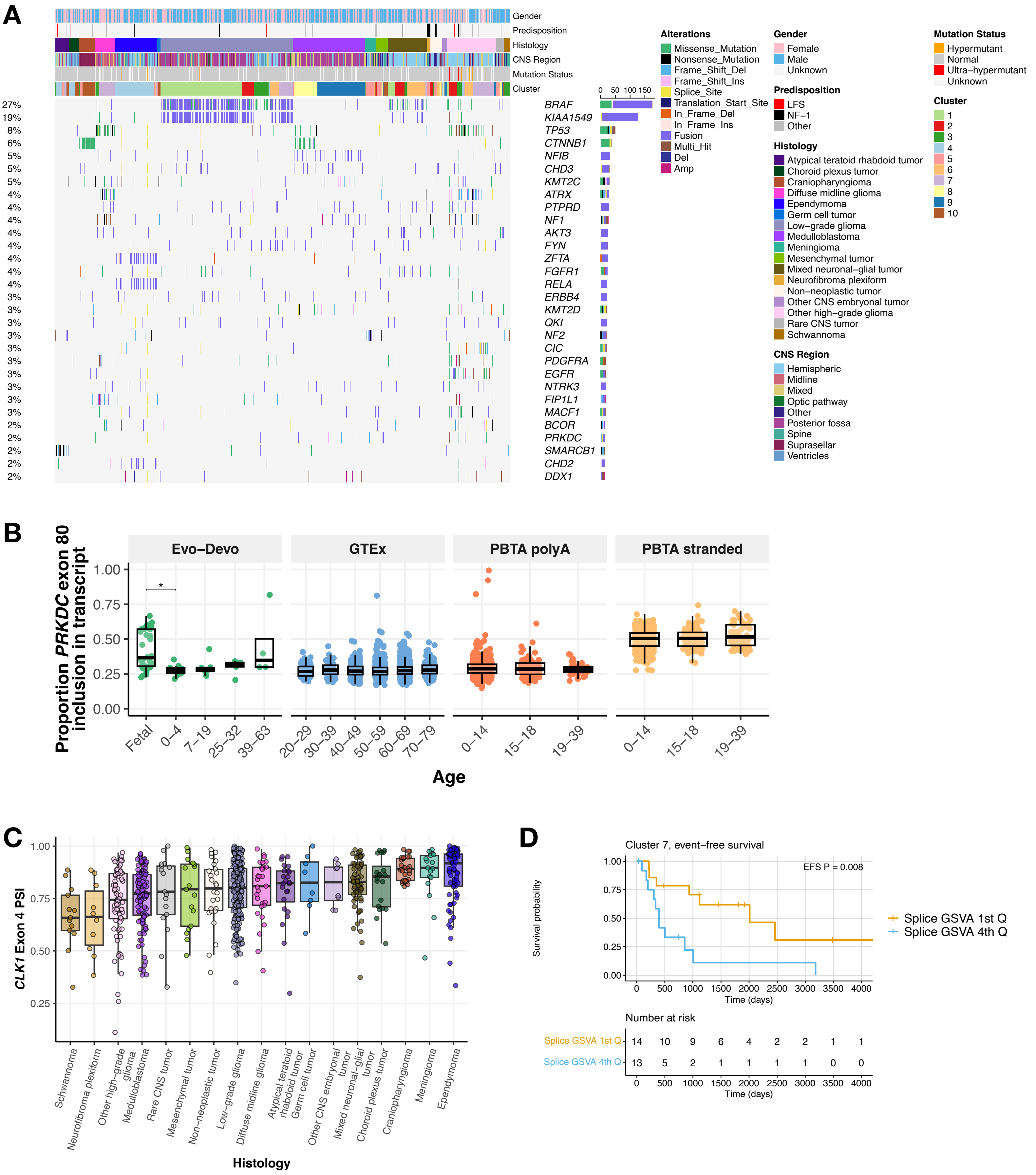

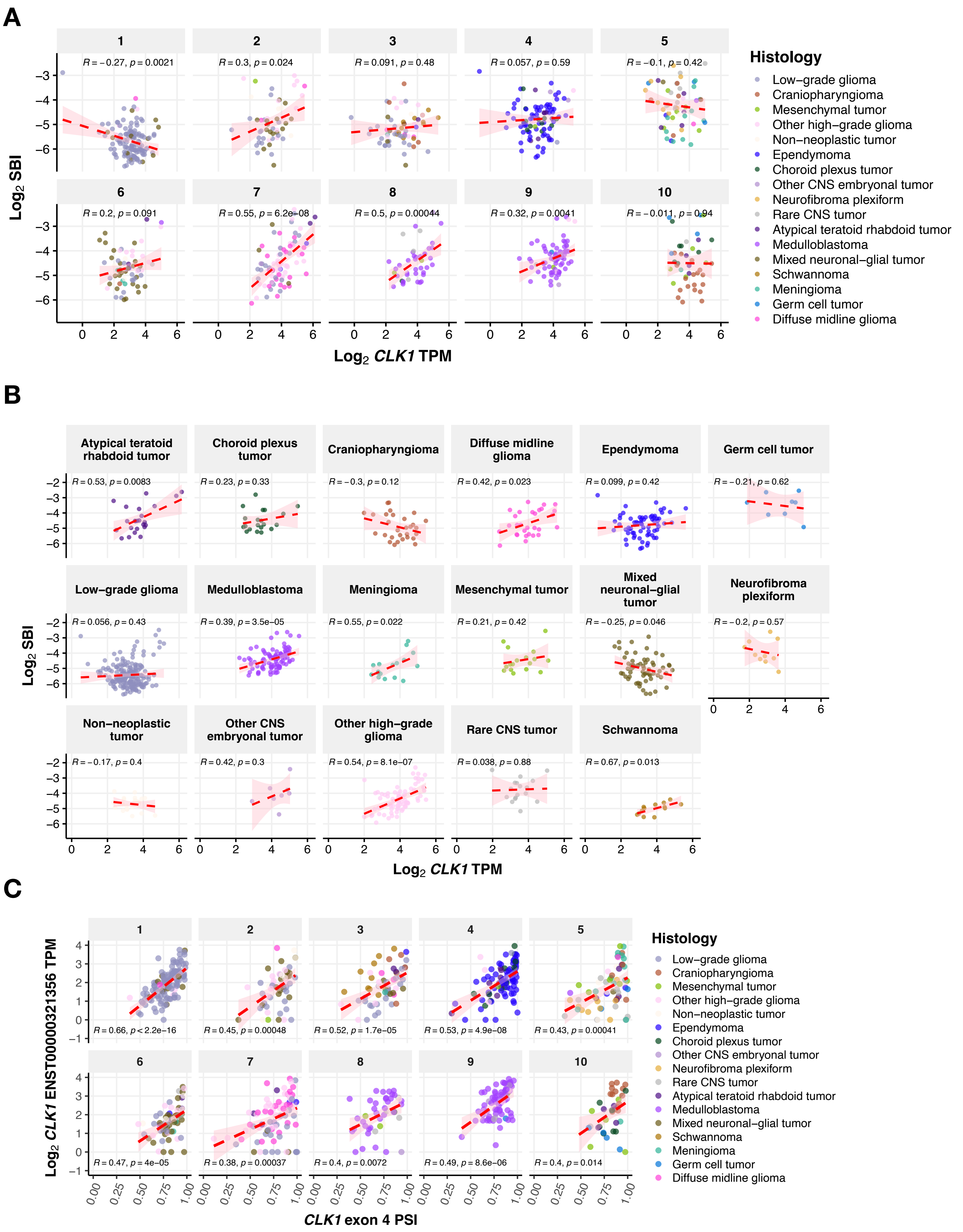

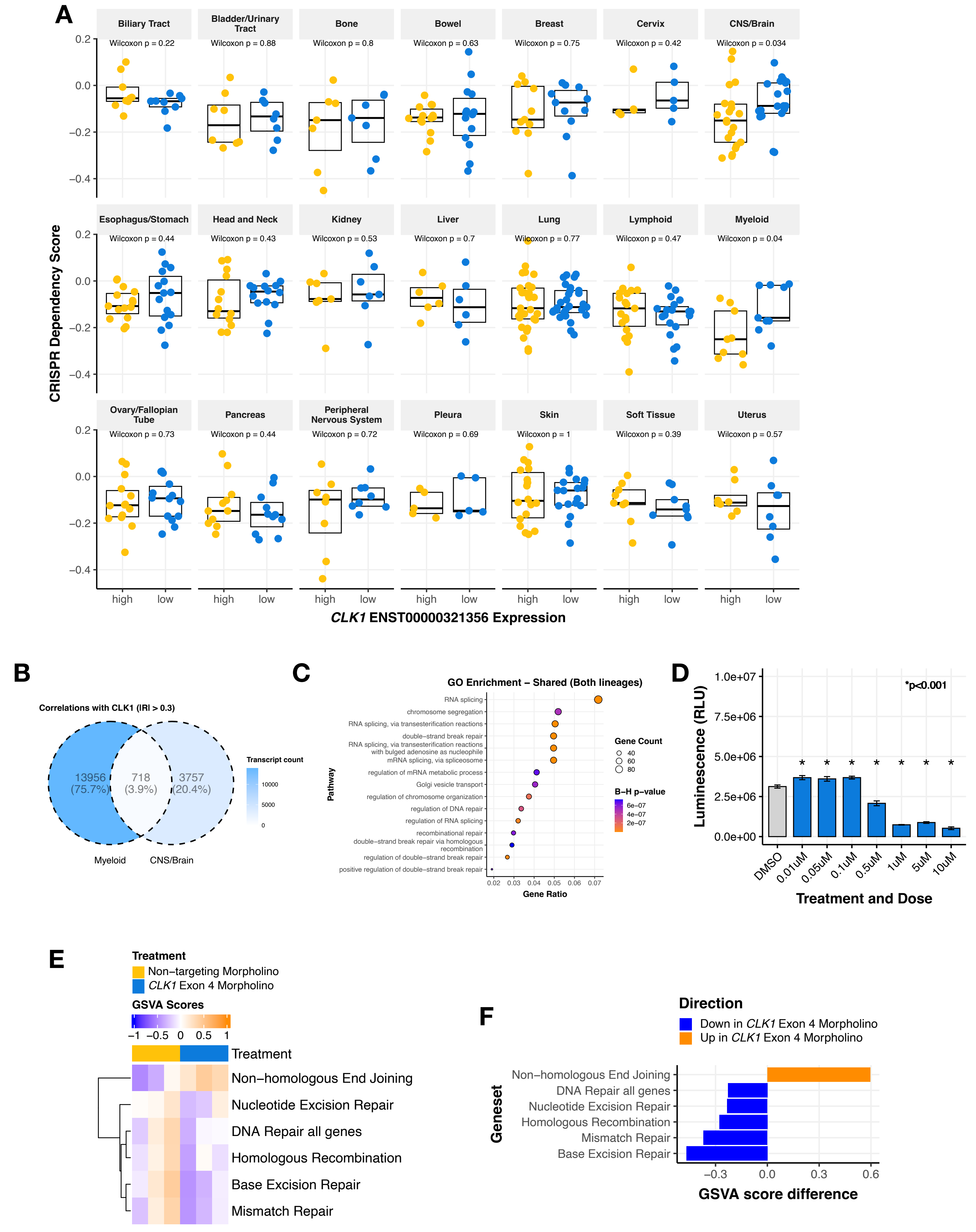
